# Dynamic Predictive Accuracy of Electrocardiographic Biomarkers of Sudden Cardiac Death within a Survival Framework: The Atherosclerosis Risk in Communities (ARIC) study

**DOI:** 10.1101/514380

**Authors:** Erick A. Perez-Alday, Aron Bender, Yin Li-Pershing, David German, Tuan Mai, Srini V. Mukundan, Christopher Hamilton, Jason Thomas, Larisa G. Tereshchenko

## Abstract

**Background:** The risk of sudden cardiac death (SCD) is known to be dynamic. However, an accuracy of a dynamic SCD prediction, and “expiration date” of ECG biomarkers is unknown. Our goal was to measure dynamic predictive accuracy of ECG biomarkers of SCD and competing outcomes.

**Methods:** Atherosclerosis Risk In Community study participants with analyzable digital ECGs were included (n=15,768; 55% female, 73% white, age 54.2±5.8 y). ECGs of 5 follow-up visits were analyzed. Global electrical heterogeneity (GEH) and traditional ECG metrics were measured. Adjudicated SCD served as the primary outcome; non-sudden cardiac death served as competing outcome. Time-dependent area under the (receiver operating characteristic) curve (AUC) analysis was performed to assess prediction accuracy of a continuous biomarker in a period of 3,6,9 months, and 1,2,3,5,10, and 15 years, using survival analysis framework.

**Results:** Over a median 24.4 y follow-up, there were 581 SCDs (incidence 1.77 (95%CI 1.63-1.92)/1,000 person-years), and 838 nonSCDs [2.55 (95%CI 2.39-2.73)]. Resting heart rate was the strongest (AUC 0.930) short-term (3-month) non-specific SCD predictor, whereas spatial peak QRS-T angle predicted specifically SCD 15 years after ECG recording (AUC 0.719). QRS duration (AUC 0.885) and QTc (AUC 0.711) short-term predicted advanced structural heart disease better than SCD. “Expiration date” for most ECG biomarkers was two years after ECG recording. GEH significantly improved reclassification of SCD risk beyond age, sex, race, diabetes, hypertension, coronary heart disease and stroke.

**Conclusion:** Short-term predictors of SCD, nonSCD, and biomarkers of long-term SCD risk differed, reflecting differences in transient vs. persistent SCD substrates.

## Introduction

Sudden cardiac death (SCD) is a major contributor to cardiovascular mortality, accounting for 40-50% of the years of potential life lost from all cardiovascular disease (CVD).^1, 2^ In the United States (US), an estimated 356,461 emergency medical services-assessed out-of-hospital sudden cardiac arrests occur annually ^1^. There remains a lack of reliable, dynamic predictors of SCD.

An electrocardiogram (ECG) can characterize the presence and properties of electrophysiological substrate of SCD. Our group recently showed that global electrical heterogeneity (GEH), as measured by five metrics [spatial QRS-T angle, spatial ventricular gradient (SVG) azimuth, elevation, and magnitude, and sum absolute QRST integral (SAI QRST)] is independently (after comprehensive adjustment for time-updated CVD events and their risk factors) associated with SCD, representing an underlying substrate of SCD ^3^. The subsequent discovery of 10 genetic loci, associated with GEH at a genome-wide significance level confirmed the presence of several underlying mechanisms behind the GEH ECG phenotype^4^. We also developed a competing risk score of SCD and showed that the addition of GEH measures to clinical risk factors significantly improves reclassification of SCD risk^3^.

The risk of SCD is known to be dynamic. However, current risk models predict SCD using baseline risk factors measured at a single point in time. An assessment of the accuracy of a dynamic prediction is therefore necessary, both to determine “an expiration date” for ECG measurements and to better understand the temporal relationship between substrate and events.

The goal of this study was to investigate the dynamic predictive accuracy of GEH and traditional ECG biomarkers of SCD within a survival framework in comparison with competing non-sudden cardiac death (nonSCD) in the Atherosclerosis Risk in Community (ARIC) study participants.

## Methods

### Study Population

The ARIC study is an ongoing prospective cohort study evaluating risk factors, progression, and outcomes of atherosclerosis in 15,792 participants (45% male, 74% white) enrolled in four US communities in 1987-1989. The ARIC study protocol and design have been previously described^5^. The study protocol was approved by institutional review boards (IRB) at each field center and all participants signed informed consent. The study was approved by the Oregon Health & Science University IRB; the study protocol conforms to the ethical guidelines of the 1975 Declaration of Helsinki.

We included all ARIC participants with available and analyzable digital ECGs (n=15,768). Participants with absent or poor quality ECGs due to noise, artifacts, or missing leads (n=24) were excluded.

Prevalent coronary heart disease (CHD) was defined as a history of myocardial infarction (MI), angina pectoris, or coronary revascularization via coronary artery bypass surgery or percutaneous coronary intervention. Prevalent MI was defined as a self-reported history of MI and/or ECG evidence of MI as defined by the Minnesota code^6^. Prevalent atrial fibrillation (AF) was defined as either a self-reported and validated history of AF or diagnosis of AF on the baseline ECG. Prevalent heart failure (HF) was defined as self-reported use of HF medication or evidence of symptomatic HF as defined by stage 3 of the Gothenburg criteria,^7^ which required the presence of specific cardiac and pulmonary symptoms in addition to medical treatment of HF. Prevalent stroke in ARIC was defined by a stroke and transient ischemic attack diagnostic algorithm, as previously described^8^. Peripheral artery disease (PAD) was defined as the ankle-brachial index ≤0.90. Details of ankle-brachial index measurement in the ARIC study have been previously described ^9^. Prevalent CVD was defined as the presence of at least one baseline prevalent condition: CHD, HF, AF, stroke, PAD, atrioventricular (AV) block II-III, atrial or ventricular pacing, or Wolff-Parkinson-White ECG phenotype.

### Definition of a primary outcome: sudden cardiac death

Follow-up of ARIC participants was previously reported^10^. To identify cases of SCD in ARIC, cases of fatal CHD that occurred by December 31, 2012 were reviewed and adjudicated by a committee of physicians in two phases, as previously described^11^. CHD deaths occurring on or before December 31, 2001 were adjudicated by 5 physicians. CHD deaths occurring between January 1, 2002 and December 31, 2012 were adjudicated by a committee of 11 physicians. Available data from death certificates, informant interviews, physician questionnaires, coroner reports, prior medical history, and hospital discharge summaries were reviewed, in addition to circumstances surrounding the event. Each event was adjudicated independently by two physicians. In cases of disagreement, a third reviewer independently reviewed the event to provide final classification. SCD was defined as a sudden pulseless condition in a previously stable individual without evidence of a non-cardiac cause of cardiac arrest if the cardiac arrest occurred out of the hospital or in the emergency room. Details of SCD adjudication are provided in the Supplement. Definite, probable, or possible SCD was included in this study as a primary outcome: the strength of available evidence determined this stratification. Witnessed SCD with an available rhythm strip of life-threatening arrhythmia, or primary emergency medical services (EMS) impression of cardiac arrest constituted definite SCD events. Probable SCD was defined as SCD with greater uncertainty either due to (i) other clinical conditions that can cloud the issue of exact cause of demise; or (ii) limited information to adjudicate an event with greater confidence. Possible SCDs were adjudicated only at the second phase of reviews (2002-2012). Unwitnessed events with limited available data but specified SCD on a death certificate, without documentation of another cause of death, were defined as possible SCD. Inter-reviewer agreement for the Phase 2 effort was 83.2% and agreement across phases was 92.5%. Participants were censored at the time of loss to follow-up or death if the cause of the death was any other than SCD.

### Competing mortality outcome: non-sudden cardiac death

Cases of fatal CHD were adjudicated as described above. Fatal CHD that did not meet criteria of SCD comprised the nonSCD outcome.

### Electrocardiogram recording and analyses

12-lead ECG was recorded according to the Atherosclerosis Risk In Communities (ARIC) study protocol and manual (version 1.0; August, 1987). The method and procedure for 12-lead ECG recording, as described in the ARIC manual, are outlined below. During the baseline examination, a standard supine 12-lead ECG was recorded after 12-hour fast followed by a light snack and at least one hour after smoking or ingestion of caffeine. The standard electrocardiograph for the ARIC study was the MAC PC by Marquette Electronics, Inc. A 12-lead resting ECG was obtained consisting of 10 seconds of each of the leads simultaneously (I, II, III, aVR, aVL, aVF, Vl-V6).

Because it is essential for the study to be able to compare baseline ECG data with subsequent records, a uniform procedure for electrode placement and skin preparation was required. The participant, stripped to the waist, was instructed to lie on the recording bed with arms relaxed at the sides. The individual was asked to avoid movements which may cause errors in marking the electrode locations. For best electrode/skin interface, the electrodes were placed on the skin at least 2-3 minutes before taking the ECG. A pen was used to mark six chest electrode positions.

The chest was wiped with a sterile alcohol prep. Left leg (LL) electrode was placed on the left ankle (inside). Right leg (RL) electrode was placed on the right ankle (inside). Left arm (LA) electrode was placed on the left wrist (inside). Right arm (RA) electrode was placed on the right wrist (inside). Electrode V1 was located in the 4^th^ intercostal space at the right sternal border, immediately to the right of the sternum. Electrode V2 was located in the 4^th^ intercostal space, immediately to the left of the sternal border. Next, E-point was located by finding the 5^th^ intercostal space, and following it horizontally to the midsternal line. Location of V6 electrode was found using the chest square. V6 was located at the same level as the E point in the midaxillary line (straight down from the center of the armpit). If breast tissue was over the V6 area, V6 was placed on top of the breast. No attempt was made to move the breast. Electrode V4 was located using a flexible ruler, as a midway between E and V6. Electrode V3 was located using a flexible ruler, midway between V2 and V4. Using a flexible ruler, electrode V5 was located midway between the locations of V4 and V6. After placing the electrodes on the skin, participant’s information was entered into MAC PC. It was required that electrodes must be on the skin for at least 3 minutes before taking the ECG. During the ECG recording, special attention was paid to the quality of recording. Quality control and technical troubleshooting procedures were in place to minimize errors (lead switch), noise, and artefacts.

Recorded 12-lead ECGs were saved in the memory of the ECG machine, and transmitted to Halifax ECG Computing Center via the phone line. Recorded 12-lead ECGs were stored in the MUSE database (GE Marquette, Milwaukee, WI). Later, MUSE database was transferred from the Halifax ECG Computing Center to Epidemiological Cardiology Research Center (EPICARE, Wake Forest University, NC). Then, stored in a MUSE database (GE Marquette, Milwaukee, WI) ECGs were exported and transferred from EPICARE to Tereshchenko laboratory at the Oregon Health & Science University (OHSU).

ECG data of all five follow-up visits were analyzed. Traditional ECG intervals were reported by the 12 SL algorithm (GE Marquette Electronics, Milwaukee, WI). QT interval was corrected for heart rate according to Bazett’s formula.

### ECG analysis: measurement of global electrical heterogeneity

We analyzed raw, digital, 10-second, 12-lead ECGs (sampling rate of 500Hz and amplitude resolution of 1μV). Origin and conduction path of each cardiac beat was adjudicated by the team of physicians (DG, AB, TM, SM, LGT), and each beat was manually labeled by investigators (CH, JAT) for subsequent automated analyses. A representative median beat was constructed for the dominant type of beat – i.e., only one type of beat was included in the development of a median beat. For development of a normal sinus median beat, sinus beats before and after PVCs and both types of noisy or distorted beats were excluded. In this study, we constructed normal sinus, junctional, supraventricular, atrial paced, and ventricular paced median beats. For consistent longitudinal analyses, we required the same type of median beat across multiple follow-up visits. For each participant, only one median beat was included in the analysis. In addition, we performed a sensitivity analysis on exclusively normal sinus median beats, which did not change the overall results (data not shown). On average, 9±2 beats were included in a median beat. The distribution of number of beats included in a median beat is shown in Figure 1. An example of a time-coherent median beat constructed in AF is shown in Figure 2.

**Figure 1.**
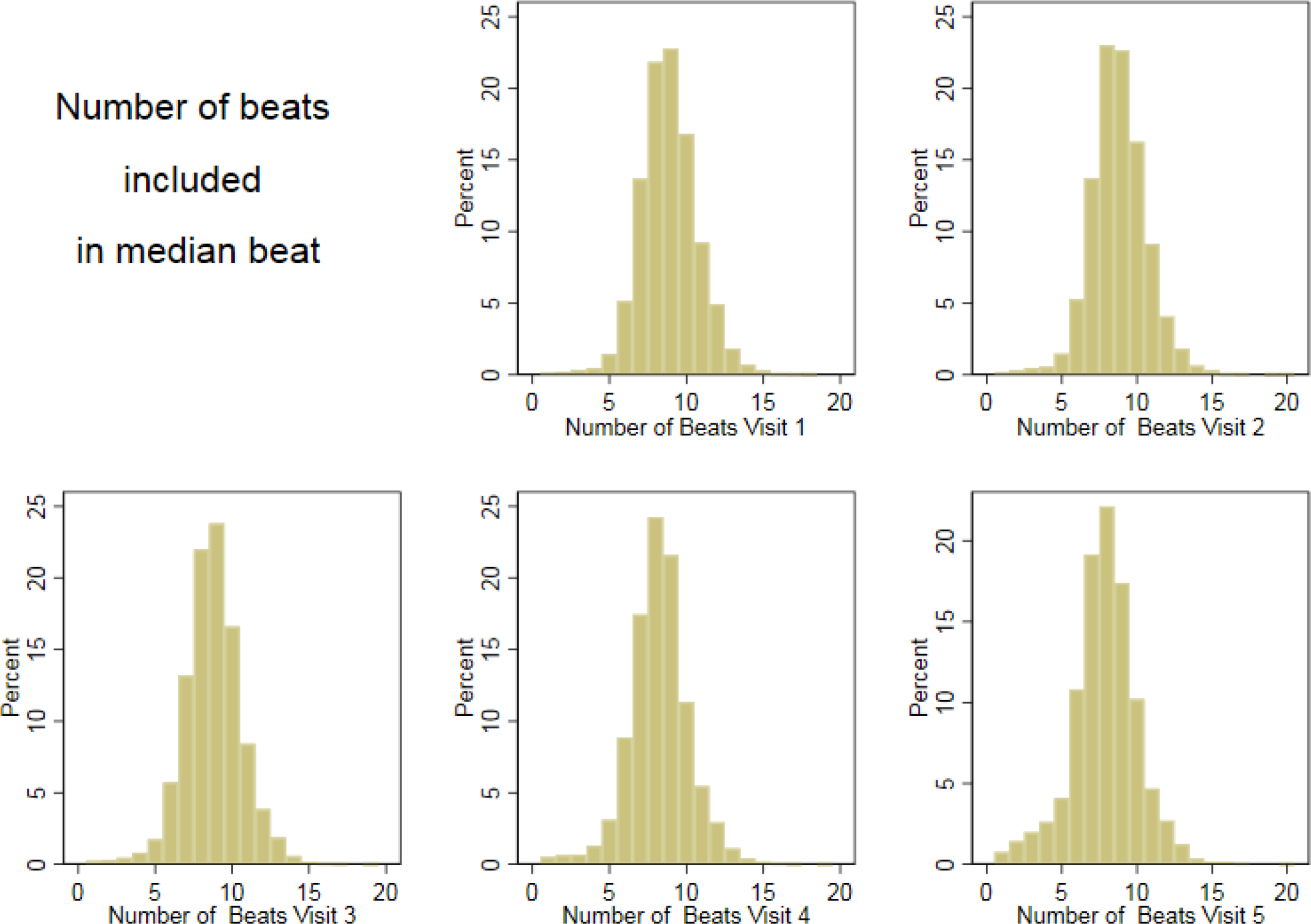
Distribution of a number of beats included in a median beat, in five study visits.

**Figure 2.**
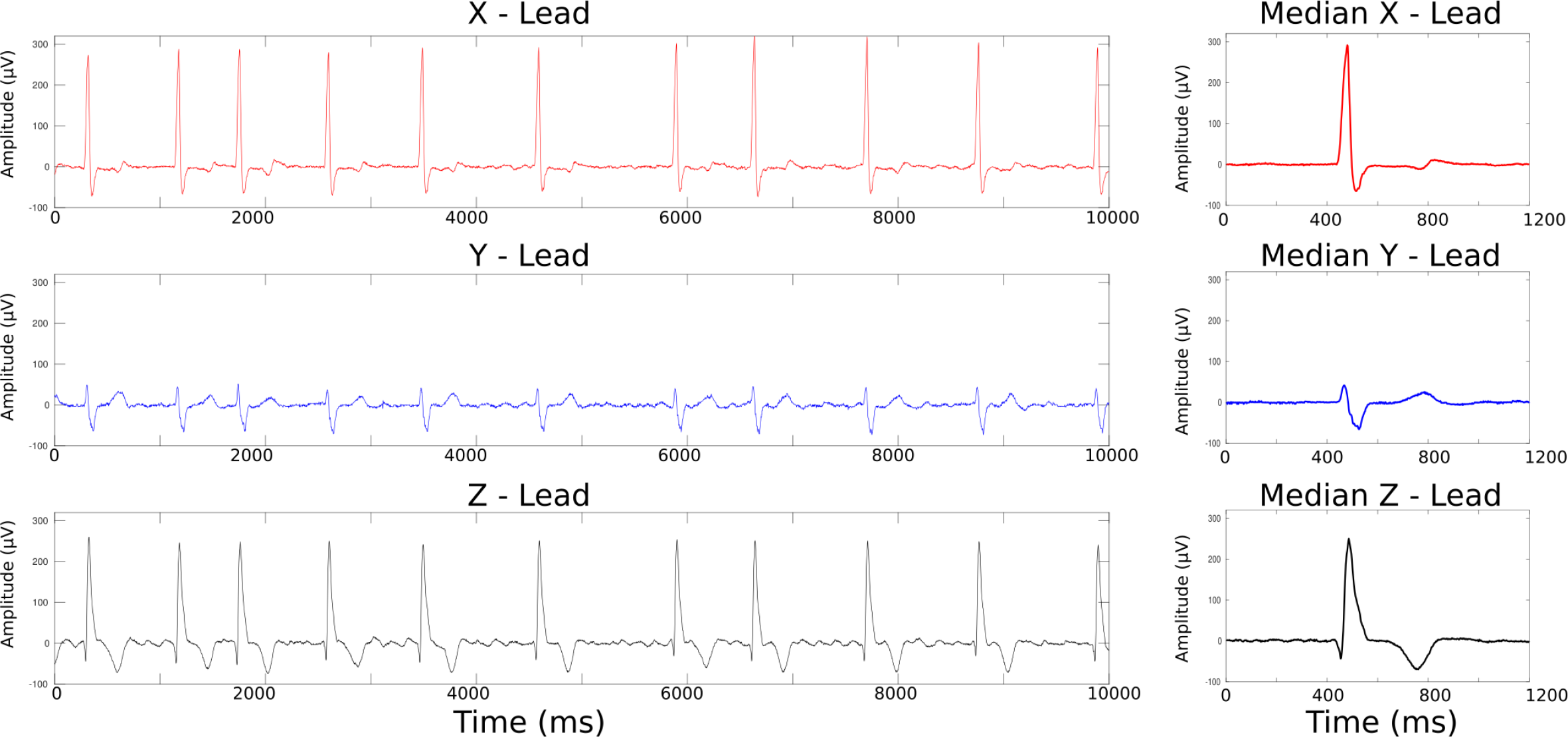
A representative example of time-coherent median beat construction in atrial fibrillation. Fibrillatory waves cancel each other in a median beat, producing clean beat, allowing accurate measurements.

GEH was measured as previously described,^3, 12^ in a time-coherent median meat with identified origin of the heart vector.^13^ We have provided the software code at Physionet (https://physionet.org/physiotools/geh/). In addition to previously reported “mean” GEH measures,^3^ in this study we measured the spatial peak vectors (Figure 3) ^12–14^.

**Figure 3.**
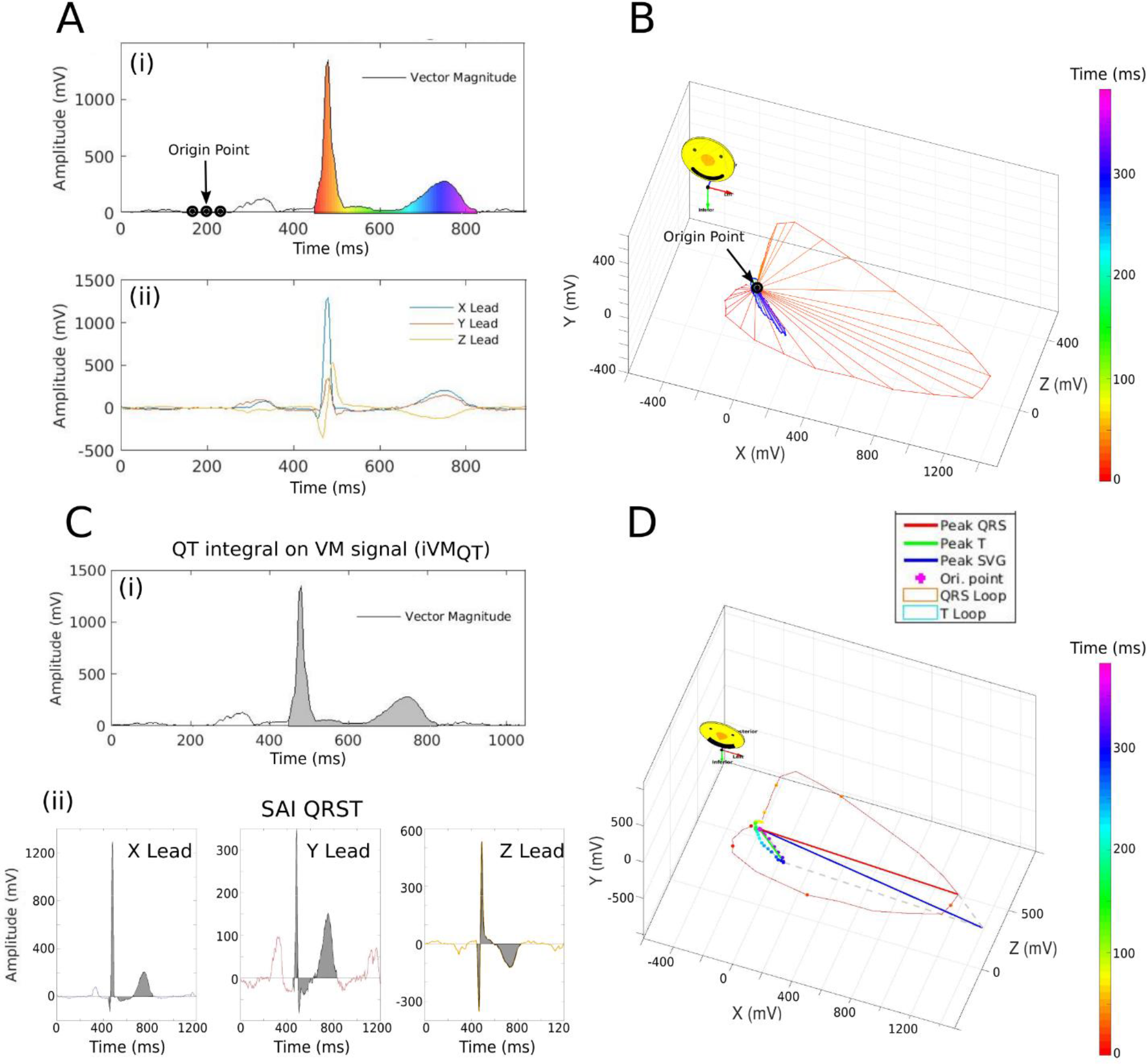
GEH measurement example of peak and area vectors, and vector magnitude (VM). (A) (i) VM plotted over time, and (ii) corresponding X, Y, and Z leads. (B) The same VM is plotted in a three-dimensional space. Color-coded progression from QRS onset (red) to the end of T (purple) is shown. (C) Measurement of (i) QT integral on VM, and (ii) sum absolute QRST integral (SAI QRST). (D). Measurement of peak spatial ventricular gradient (SVG) vector magnitude, azimuth, and elevation.

### ECG analysis: measurement of global electrical heterogeneity

At OHSU, GE Magellan research utility (GE Marquette, Milwaukee, WI) was used to obtain raw digital 12-lead ECG signal (sampling rate 500 Hz; amplitude resolution 1μV), as well as fiducial points (QRS onset and offset, T offset) that were used for measurement of “area” vectors and sum absolute QRST integral (SAI QRST).

In this study, we calculated both types of spatial ventricular gradient (SVG) vectors, and spatial QRS-T angle: “area” and “peak”, as we previously described. ^12^ SAI QRST was measured as the arithmetic sum of areas under the QRST curve on X, Y, and Z leads, as previously described.^3^

First, we transformed 12-lead ECG in orthogonal XYZ ECG, using Kors transformation.^15^ Next, we constructed time-coherent median beat, and detected origin of the heart vector, using our novel approach, as recently described.^13^ Then, we performed calculations of GEH metrics, using the following equations.

#### Spatial QRS-T angles

Spatial peak QRS-T angle was calculated as the 3-dimensional angle between the spatial peak QRS vector and the spatial peak T vector:

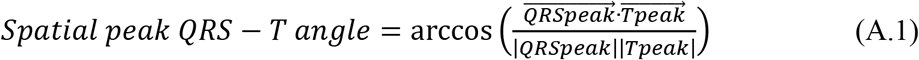

Spatial area QRS-T angle was calculated as the 3-dimensional angle between the spatial area QRS vector and the spatial area T vector:

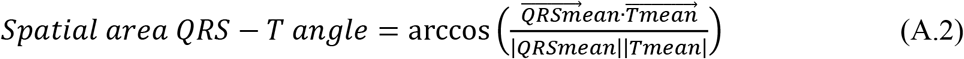

#### Spatial ventricular gradient vectors

Magnitude and direction of spatial area (Wilson’s) and peak SVG vectors were measured.

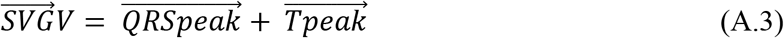

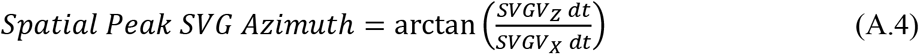

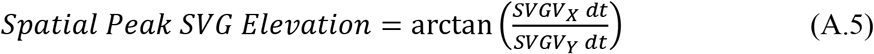

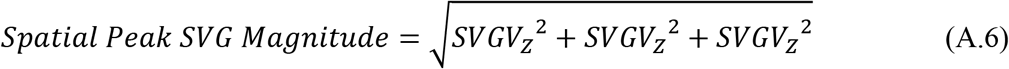

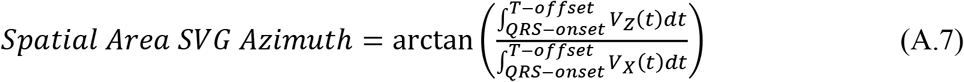

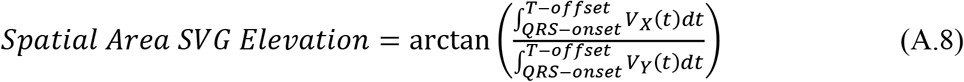

Wilson’s (area) SVG magnitude was also calculated:

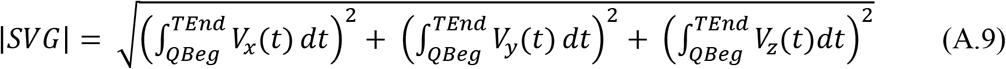

We provided MATLAB (MathWorks, Natick, MA, USA) code as an open source at: https://github.com/Tereshchenkolab/Global-Electrical-Heterogeneity https://github.com/Tereshchenkolab/Origin

### Statistical analysis

Time-dependent area under the (receiver operating characteristic) curve (AUC) analysis was performed to assess predictive accuracy of a continuous biomarker in a period of 3, 6, 9 months, and 1,2,3,5,10, and 15 years, using an unadjusted survival analysis framework approach^16, 17^. We used the nearest neighbor estimator which allows the censoring to depend on the marker and is therefore realistic. The percentage of observations included in each neighborhood was defined by equation 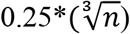, where *n* is the number of observations. All available five visits’ ECG data were included in time-dependent AUC analysis.^3^ Equality of AUC areas for different ECG and VCG biomarkers of SCD and competing mortality outcomes was compared using the Wald test.^18^ The best thresholds were selected using Liu’s optimal survival time-dependent cut-point on the prognostic biomarker.^19^We summarized clinical characteristics of study participants with a SCD outcome within the first 3 months, 3-6 months, 6 months-1 year, 1-2 years, 2-5 years, and more than 5 years after ECG recording in a longitudinal dataset, reporting between-participant standard deviation (SD) for continuous variables, and between-participant frequencies for categorical variables.

We then assessed whether the addition of traditional ECG metrics (heart rate, QRS, QTc) and GEH metrics to our previously identified clinical risk factors of SCD^3^ (age, sex, race, diabetes, hypertension, CHD, and stroke) resulted in better predictive accuracy for SCD and nonSCD within the first 3 months, 3-6 months, 6 months-1 year, 1-2 years, 2-5 years, and more than 5 years after ECG recording. We calculated absolute integrated discrimination improvement (IDI), and net reclassification improvement (NRI) using multivariate logistic regression.^20, 21^ IDI estimates improvement in average sensitivity and specificity. We estimated category-free NRI and two-category NRI for events, defining the high risk category as a ≥25% risk of SCD/nonSCD within the first 3 months, 3-6 months, and 6 months-1 year after ECG recording. The high risk category for events occurring 1-2 years, 2-5 years, and more than 5 years after ECG recording was defined as ≥ 10% risk of SCD/nonSCD.

Also, as resting heart rate is a known predictor of cardiovascular death, we assessed whether the addition of GEH metrics to heart rate improves the predictive accuracy for SCD and nonSCD.

Statistical analysis was performed using STATA MP 15.1 (StataCorp LP, College Station, TX, USA). A P-value of < 0.05 was considered significant.

## Results

### Study population

Clinical characteristics of the study population are shown in Table 1. Approximately half of the study participants were female and 73% were white. Average traditional ECG parameters were normal. Over a median follow-up of 24.4 years, there were 581 SCDs (incidence 1.77 (95%CI 1.63-1.92) per 1,000 person-years), and 838 nonSCDs (incidence 2.55 (95%CI 2.39-2.73) per 1,000 person-years).

**Table 1.**
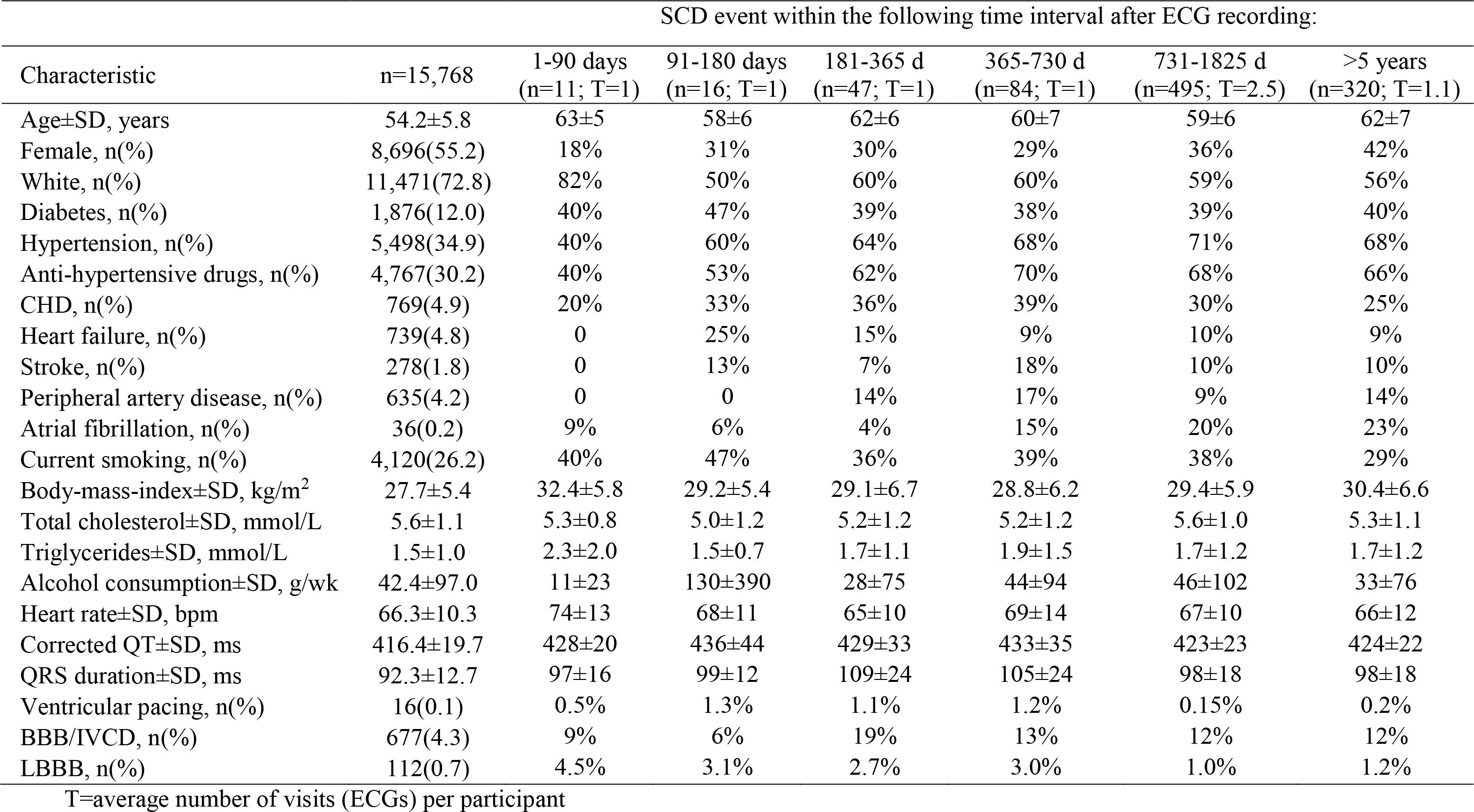
Clinical characteristics of study population.

SCD victims who died within the first three months after ECG recording were more likely to be CVD-free white males with fewer prevalent CVD risk factors. In contrast, SCD victims who died more than five years after ECG recording had nearly equal probabilities of being male or female, white or non-white (Table 1).

### Time-dependent AUC analyses of SCD and competing outcomes

All ECG biomarkers had pronounced differences in the dynamic accuracy of both mortality outcomes prediction [(Figure 4) and (Supplemental Figures 2–25)]. Overall, ECG measures were mostly predictive of outcomes within first two years after ECG recording.

**Figure 4.**
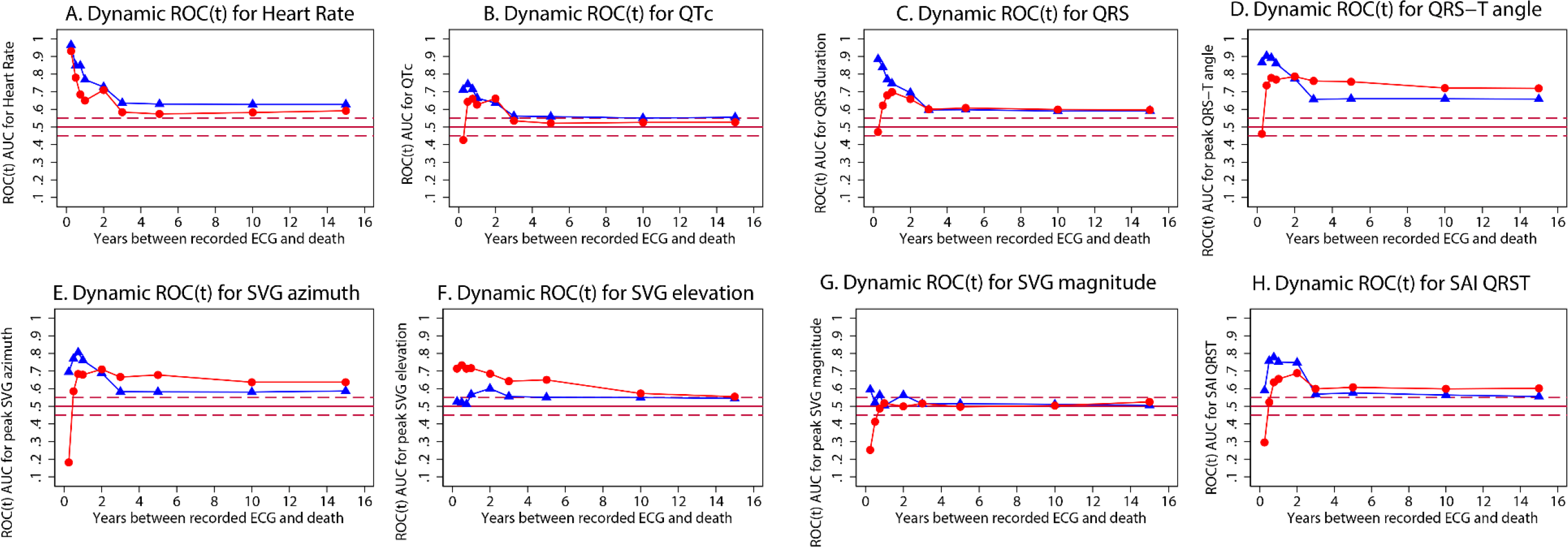
Figure 2. Time-dependent AUC for prediction of SCD (red circles), and nonSCD (blue triangles) for windows of prediction 3, 6, 9 months, 1, 2, 3, 5, 10, 15 years for (A) heart rate, (B) QTc interval, (C) QRS duration, (D) spatial peak QRS-T angle, (E) peak SVG azimuth, (F) peak SVG elevation, (G) peak SVG magnitude, (H) SAI QRST measured at visits 1, 2, 3, 4, and 5.

As expected, resting heart rate was a non-specific predictor: it similarly predicted both mortality outcomes (Figures 4A and Supplemental Figures 2–3). QTc also performed equally well in predicting SCD and nonSCD within two years after ECG recording only (Figures 4B and Supplemental Figures 4–5). Unlike QTc, QRS duration remained a weak predictor of cardiovascular mortality long-term (Figures 4C and Supplemental Figures 6–7). Neither heart rate nor QTc improved risk reclassification beyond traditional clinical risk factors of SCD (Table 2), whereas QRS duration improved reclassification of SCD risk for events occurring 2-5 years after ECG recording.

**Table 2.**
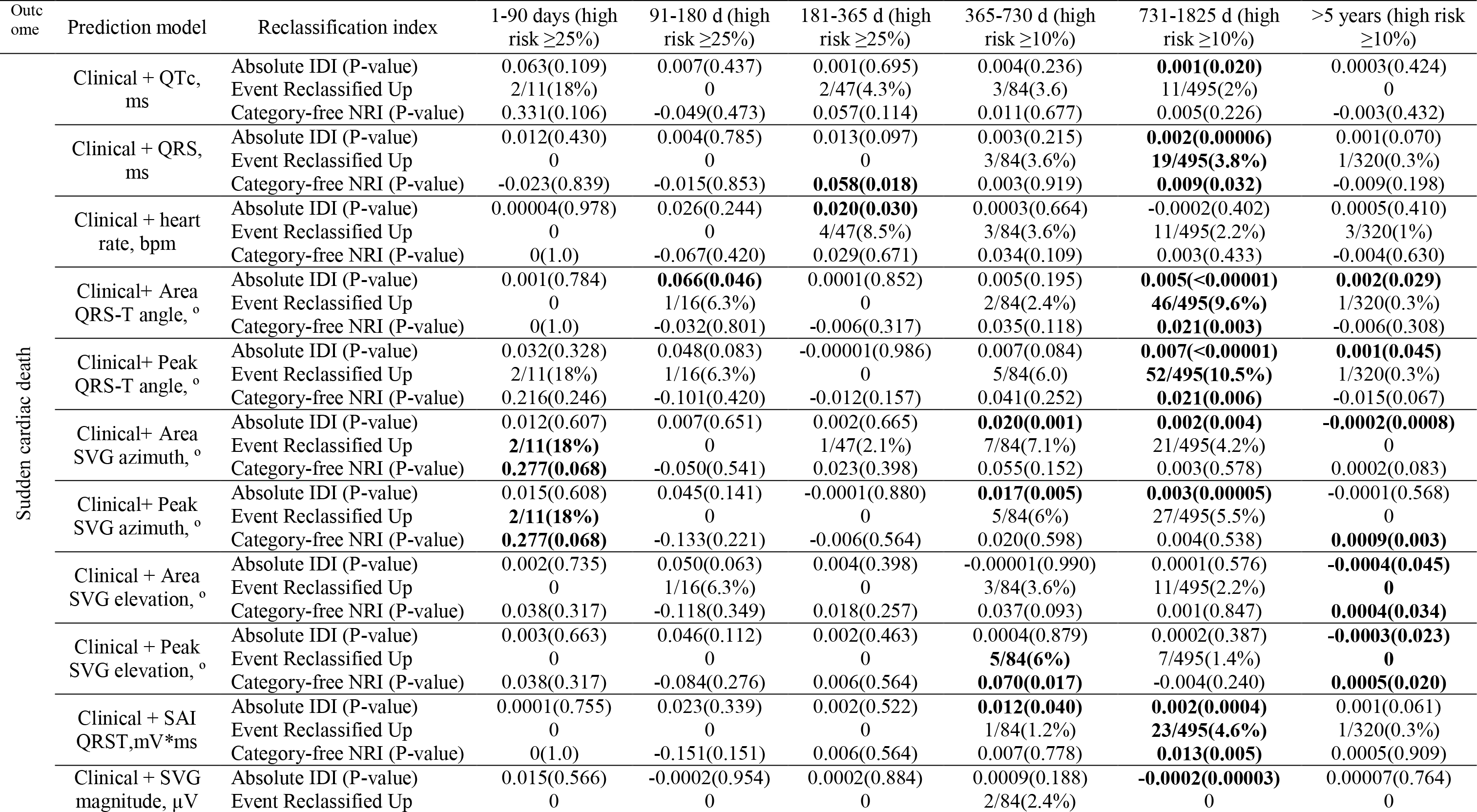
Integrated Discrimination Improvement (IDI) and Net Reclassification Improvement (NRI) for ECG metrics added to clinical predictors of SCD and nonSCD outcomes (age, sex, race, coronary heart disease, stroke, hypertension, diabetes)

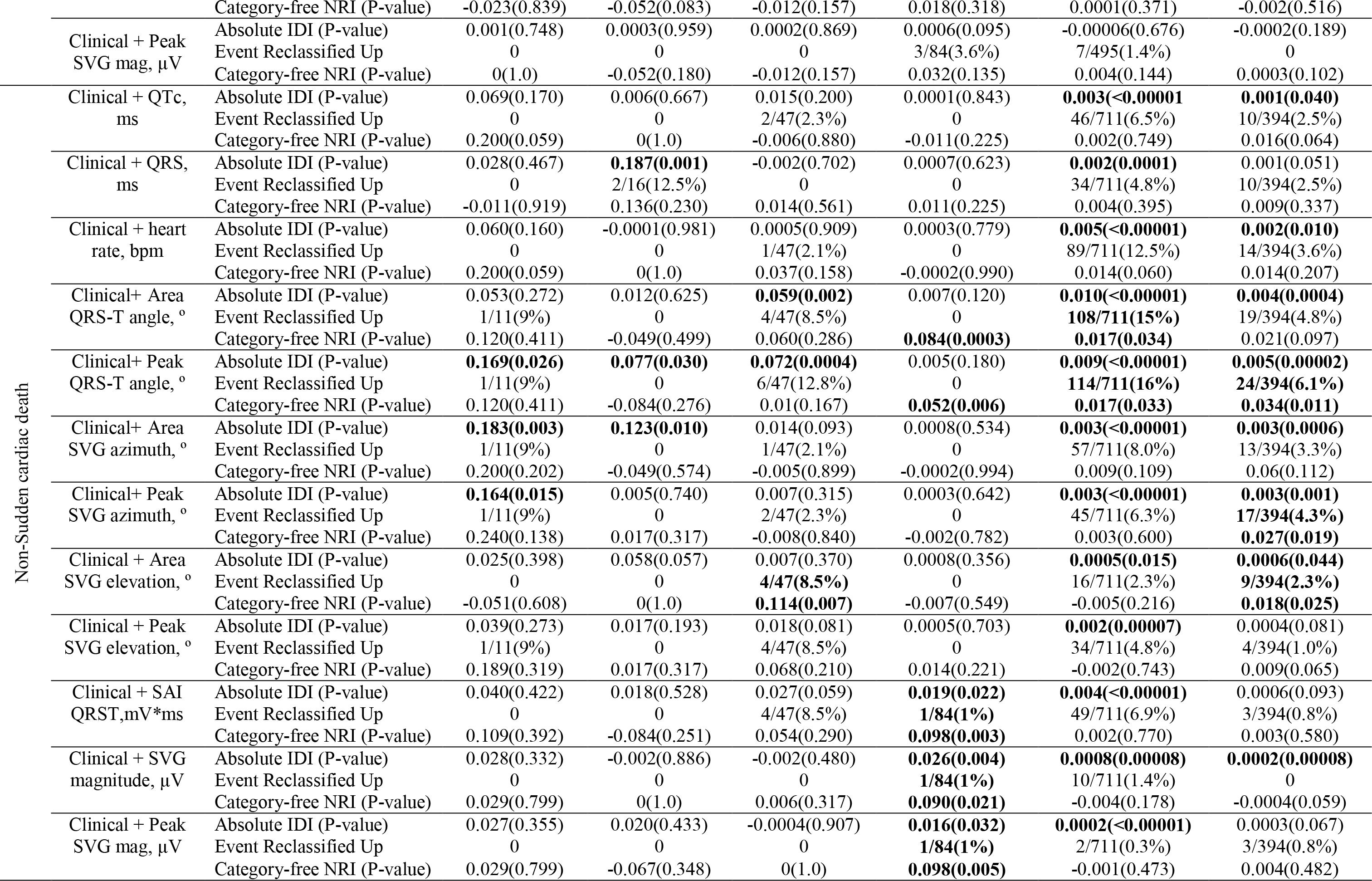

Individual GEH metrics outperformed individual traditional metrics for prediction of SCD but not for prediction of nonSCD. Spatial QRS-T angle (Figures 4D and Supplemental Figures 8–11) was the strongest long-term predictor of SCD, consistently maintaining an AUC> 0.7 across time points. SVG azimuth (Figures 4E and Supplemental Figures 12–15) also was a significantly stronger predictor of SCD as compared to nonSCD, up to 15 years after ECG recording. SVG elevation (Figures 4F and Supplemental Figures 16–19) specifically predicted SCD up to 5 years after ECG recording. SVG magnitude (Figures 4G and Supplemental Figures 20–23) differentially predicted mortality outcomes only within the first 3 months after ECG recoding but not thereafter. Except for differences in the prediction of SCD and nonSCD within the first 3 months after ECG recording, SAI QRST similarly predicted long-term SCD and nonSCD (Figures 4H and Supplemental Figures 24–25). All GEH metrics significantly improved reclassification of both SCD and nonSCD beyond clinical risk factors for events occurring at least 1 year after ECG recording (Table 2). Of note, GEH improved sensitivity in the first 1-5 years after ECG recording, whereas long-term (>5 years), GEH improved the specificity of prediction.

### Comparison of short-term and long-term predictors of SCD and competing nonSCD

For both outcomes, short-term and long-term predictors were significantly different from each other. Few ECG biomarkers predicted SCD occurring within 3 months after ECG recording (Figure 5). Resting heart rate had the best predictive accuracy in the short-term (within 3 months). However, heart rate did not improve reclassification of risk beyond clinical risk factors (Table 2). Short-term addition of GEH metrics to heart rate did not improve reclassification of the risk (Table 3). Starting in 3 months after ECG recording, all other ECG metrics improved reclassification of the risk beyond heart rate.

**Figure 5.**
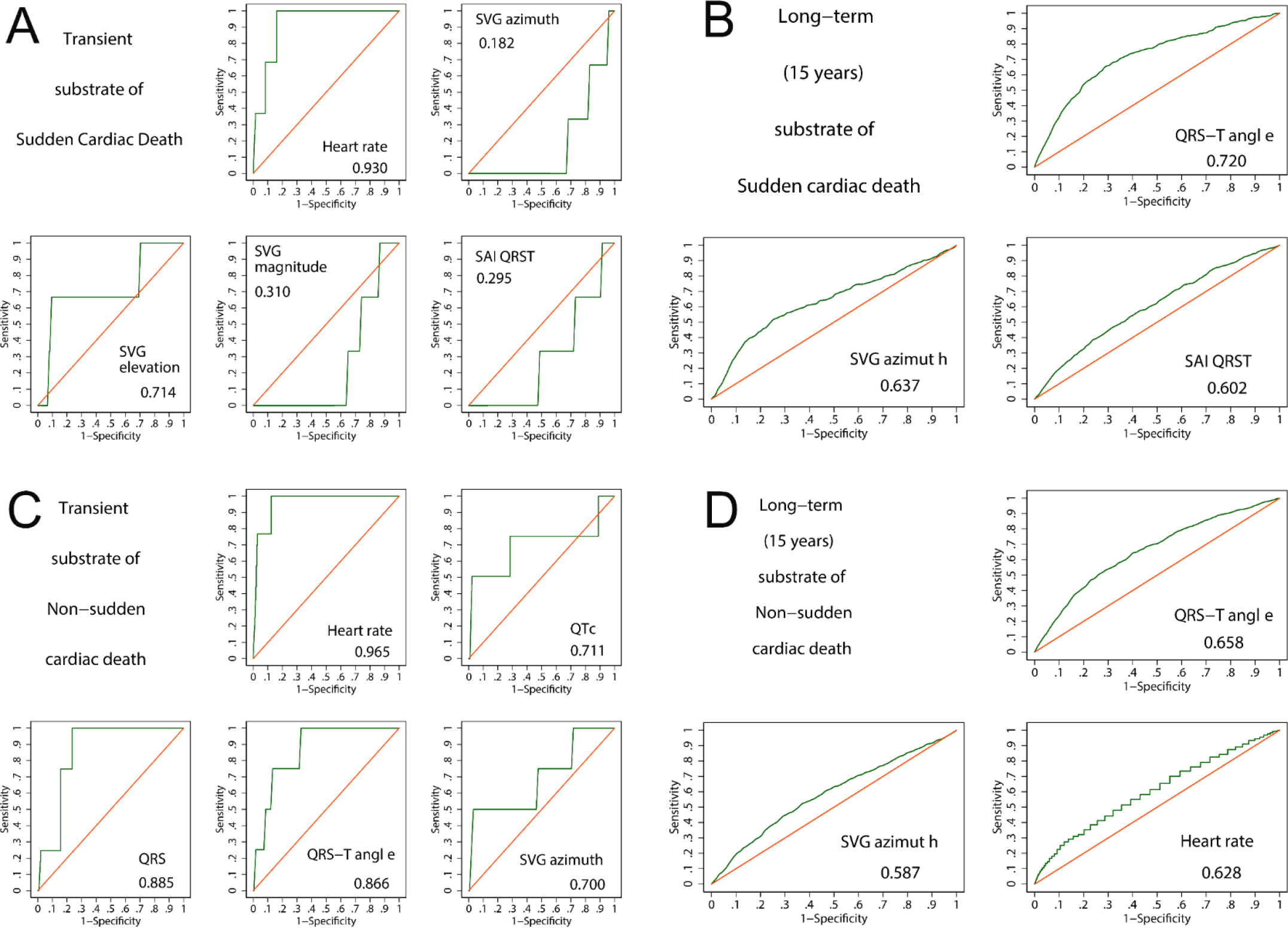
Comparison of transient (3-month) vs. long-term persistent (15-year) substrates of SCD vs. nonSCD. Time-dependent ROC curves for prediction of (A) SCD within 3 months after ECG recording, (B) SCD within 15 years after ECG recording, (C) nonSCD within 3 months after ECG recording, (D) nonSCD within 15 years after ECG recording.

**Table 3.** Absolute Integrated Discrimination Improvement for GEH metrics added to resting heart rate (HR) for prediction of sudden cardiac death and non-sudden cardiac death outcomes.

| Outcome | Prediction model | 1-90 days<br>(high risk<br>≥25%) | 91-180 d (high<br>risk ≥5%) | 181-365 d<br>(high risk<br>≥5%) | 365-730 d (high<br>risk ≥5%) | 731-1825 d (high<br>risk ≥5%) | >5 years (high<br>risk ≥5%) |
| --- | --- | --- | --- | --- | --- | --- | --- |
|  |  | Absolute Integrated Discrimination Improvement (P-value) |  |  |  |  |  |
| Sudden cardiac death | Heart rate + QTc, ms | 0.002(0.392) | 0.003(0.243) | 0.0005(0.561) | 0.002(0.140) | **0.002(<0.00001)** | 0.0003(0.128) |
|  | Heart rate + QRS, ms | 0.001(0.328) | 0.0006(0.316) | **0.008(0.023)** | **0.008(0.010)** | **0.007(<0.00001)** | **0.003(0.00001)** |
|  | HR+ Area QRS-T angle, ° | 0.004(0.175) | **0.021(0.001)** | **0.005(0.043)** | **0.011(0.0004)** | **0.013(<0.00001)** | **0.005(<0.00001)** |
|  | HR+ Peak QRS-T angle, ° | 0.001(0.195) | **0.034(0.002)** | **0.018(0.0006)** | **0.018(<0.00001)** | **0.017(<0.00001)** | **0.006(<0.00001)** |
|  | HR+ Area SVG azimuth, ° | 0.010(0.190) | 0.003(0.159) | 0.004(0.104) | **0.013(0.0007)** | **0.002(<0.00001)** | 0.0002(0.117) |
|  | HR+ Peak SVG azimuth, ° | 0.021(0.090) | **0.009(0.046)** | 0.004(0.073) | **0.011(0.002)** | **0.004(<0.00001)** | **0.001(0.004)** |
|  | HR + Area SVG elevation, ° | 0.0002(0.928) | 0.002(0.191) | 0.002(0.252) | 0.0003(0.272) | **0.004(<0.00001)** | **0.002(0.0004)** |
|  | HR + Peak SVG elevation, ° | 0.0001(0.882) | 0.002(0.257) | 0.002(0.336) | 0.0004(0.379) | **0.005(<0.0001)** | **0.002(0.001)** |
|  | HR + SAI QRST,mV*ms | 0.001(0.227) | **0.005(0.046)** | **0.017(0.037)** | **0.019(0.002)** | **0.006(<0.00001)** | **0.003(0.00007)** |
|  | HR+SVG magnitude, μV | 0.018(0.336) | 0.011(0.153) | **0.016(0.008)** | **0.008(0.039)** | **0.0003(0.0009)** | **0.0006(0.022)** |
|  | HR+Peak SVG mag, μV | 0.001(0.548) | 0.005(0.212) | 0.005(0.084) | 0.002(0.163) | 0(0.591) | 0.0002(0.199) |
| Non-Sudden cardiac death | Heart rate + QTc, ms | 0.029(0.273) | 0.008(0.498) | 0.011(0.018) | 0.0005(0.282) | **0.003(<0.0001)** | 0.0004(0.058) |
|  | Heart rate + QRS, ms | 0.025(0.158) | **0.009(0.016)** | 0.006(0.087) | **0.006(0.013)** | **0.006(<0.0001)** | **0.003(0.00001)** |
|  | HR+ Area QRS-T angle, ° | 0.023(0.278) | **0.034(0.002)** | **0.045(<0.0001)** | **0.010(0.0004)** | **0.017(<0.0001)** | **0.008(<0.0001)** |
|  | HR+ Peak QRS-T angle, ° | 0.053(0.091) | **0.059(0.0002)** | **0.074(<0.0001)** | **0.026(<0.0001)** | **0.019(<0.0001)** | **0.010(<0.0001)** |
|  | HR+ Area SVG azimuth, ° | 0.017(0.367) | **0.013(0.010)** | **0.026(0.0005)** | **0.005(0.014)** | **0.003(<0.0001)** | **0.003(<0.0001)** |
|  | HR+ Peak SVG azimuth, ° | 0.025(0.305) | 0.004(0.413) | **0.018(0.004)** | 0.002(0.149) | **0.004(<0.0001)** | **0.004(<0.0001)** |
|  | HR + Area SVG elevation, ° | 0.001(0.801) | 0.005(0.253) | **0.010(0.028)** | **0.0003(0.607)** | **0.004(<0.0001)** | **0.002(0.0001)** |
|  | HR + Peak SVG elevation, ° | 0.005(0.568) | 0.004(0.339) | **0.014(0.012)** | 0.0006(0.450) | **0.007(<0.0001)** | **0.002(0.0001)** |
|  | HR + SAI QRST,mV*ms | 0.057(0.160) | **0.013(0.006)** | **0.042(0.0006)** | **0.026(0.0001)** | **0.008(<0.0001)** | **0.002(0.0002)** |
|  | HR+SVG magnitude, μV | 0.036(0.131) | 0.014(0.064) | **0.025(0.014)** | **0.036(0.0001)** | **0.0006(0.0001)** | 0.00002(0.459) |
|  | HR+Peak SVG mag, μV | 0.017(0.178) | 0.003(0.258) | **0.011(0.046)** | **0.014(0.004)** | 0.0001(0.064) | 0.0001(0.139) |

Remarkably, short-term GEH predictors of SCD and nonSCD were dramatically different. Small SVG magnitude, small SAI QRST, and upward-forward [towards the right ventricular outflow tract (RVOT)] – directed SVG vector predicted short-term SCD. In contrast, backward [towards the left ventricle (LV)] – directed SVG vector predicted nonSCD. QTc, QRS duration, and QRS-T angle predicted nonSCD, but not SCD (Figure 5 A,C). Within the first 3 months after ECG recording, only SVG azimuth improved reclassification of the risk beyond traditional clinical risk factors (Table 2).

In contrast, long-term predictors of SCD and nonSCD had many similarities. Spatial QRS-T angle and backward [towards LV] – directed SVG azimuth predicted both SCD and nonSCD. However, QRS-T angle and SVG azimuth were significantly stronger predictors of SCD than nonSCD (P<0.05). SAI QRST predicted SCD whereas resting heart rate predicted nonSCD (Figure 5 B,D). The best threshold was relatively stable over time (Table 4). Long-term GEH metrics significantly improved reclassification of risk beyond clinical risk factors (Table 2).

**Table 4.**
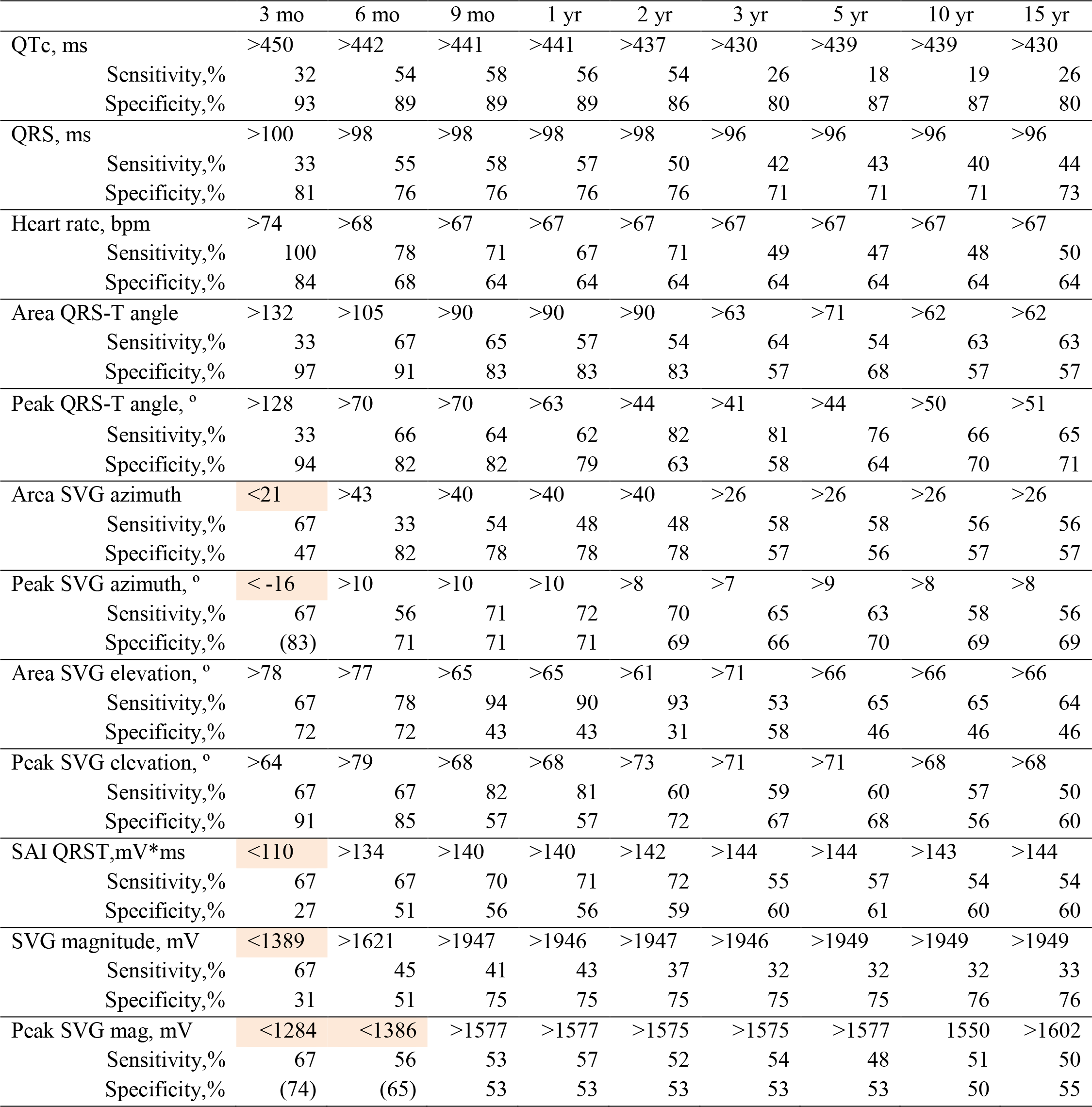
Time-dependent thresholds of ECG measures for SCD prediction, and their sensitivity/specificity.

SCD predictive accuracy of peak-vector based spatial QRS-T angle and SVG azimuth (but not SVG elevation) was significantly better, as compared to the predictive accuracy of respective area-based metrics (Table 5; Figure 6).

**Figure 6.**
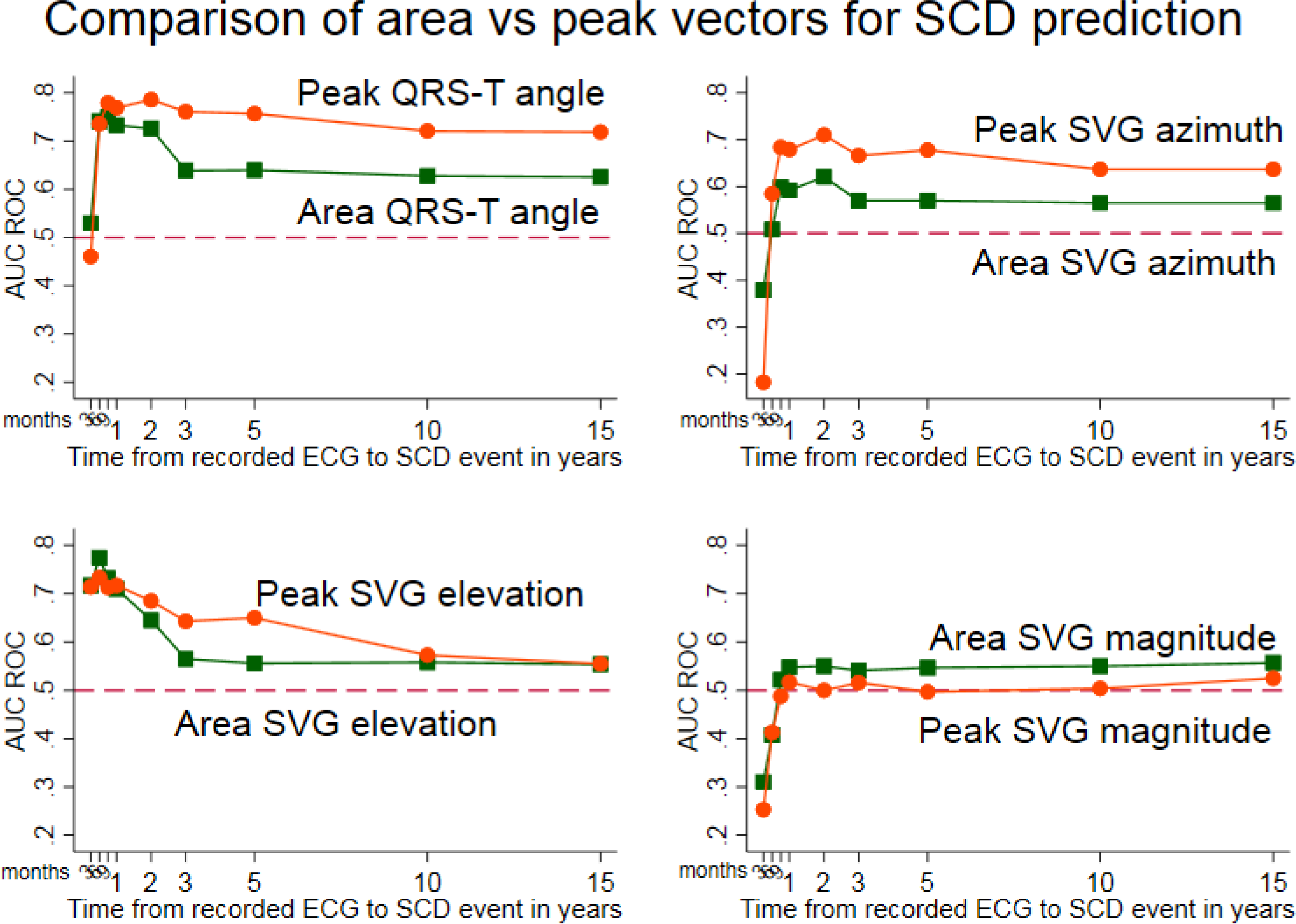
Comparison of time-dependent AUC for windows of SCD prediction 3, 6, 9 months, 1, 2, 3, 5, 10, 15 years, for peak-based SVG vector measurements (orange circles) vs. area-based SVG vector measurements (green squares).

**Table 5.** Comparison of spatial vector-based vs. area-based GEH measurements on predictive accuracy for SCD prediction, measured by AUC with 95% Confidence Interval.

| Measurement | Peak vectors AUC (95% CI) | Areas vectors AUC (95% CI) | P-value |
| --- | --- | --- | --- |
| QRS-T angle | 0.676(0.652-0.699) | 0.638(0.64-0.663) | 0.0001 |
| SVG azimuth | 0.571(0.545-0.597) | 0.542(0.516-0.567) | 0.007 |
| SVG elevation | 0.608(0.584-0.632) | 0.607(0.584-0.632) | 0.902 |
| SVG magnitude | 0.498(0.473-0.522) | 0.512(0.490-0.540) | 0.001 |

## Discussion

In this study, we described the dynamic predictive accuracy of ECG and VCG biomarkers of competing mortality outcomes: SCD and nonSCD within a survival framework. All ECG biomarkers more accurately predicted events that occurred within first 2 years after ECG recording, as compared to events that occurred later. Therefore, future dynamic risk scores of SCD should consider inclusion of every two-year-updated ECG biomarkers. Within the identified dynamic predictors of SCD, there was a distinction between markers predicting short-term events (within 3 months) and markers predicting more intermediate-and long-term events. This may represent the difference between markers heralding SCD (triggers or transient substrates) versus markers identifying persistent substrate. Spatial peak QRS-T angle was the strongest long-term (15 years after ECG recording) SCD-specific predictor, and, therefore, could be considered for a life-long SCD prediction. As expected, transient substrate of nonSCD (describing structural heart disease substrate) was characterized by wide QRS-T angle, SVG vector pointing towards LV, wide QRS, prolonged QTc, and increased heart rate. Transient substrate of SCD was uniquely characterized by SVG vector pointing towards RVOT and small SAI QRST and SVG magnitude. Dynamic predictive accuracy and knowledge of an “expiration date” of ECG and VCG biomarkers of SCD should be taken into account for development of dynamic and life-long prediction of SCD and nonSCD. Importantly, the addition of GEH metrics (but not QTc or QRS duration) to known demographic and clinical risk factors (age, sex race, CHD, stroke, diabetes, and hypertension) significantly improved reclassification of risk, supporting inclusion of GEH metrics into dynamic risk scores for SCD.

### Triggers, or transient substrate of SCD event

A SCD event represents a “perfect storm”, requiring both susceptible anatomical/functional substrate and a trigger/transient initiating event.^22^ Short-term predictors of SCD in our study reflect possible SCD triggers. As expected, resting heart rate was the strongest non-specific predictor of SCD within three months before the SCD event. Sinus tachycardia is a marker of increased sympathetic tone, ^23, 24^ a well-recognized trigger of SCD.

The SVG vector direction predicting short-term SCD risk differed from SVG vector direction of the intermediate-and long-term risk of SCD. Short-term risk was uniquely predicted by an SVG vector pointing upward and forward toward the RVOT, suggesting that a short total recovery time in the outflow tracts (as the SVG vector points towards an area with the shortest total refractory time)^25^ may represent an SCD trigger. SVG azimuth was the only ECG metric which improved reclassification of the risk beyond known clinical and demographic risk factors. Indeed, it is known that “malignant” idiopathic ventricular fibrillation and polymorphic ventricular tachycardia can be triggered by ventricular ectopy arising from the outflow tracts.^26^ Early cardiac development affects the generation of electrophysiological heterogeneities in the adult heart.^27^ There may be a genetic basis for this phenomenon as GEH-associated genetic polymorphisms indicated the involvement of *HAND1* and *TBX3* genes,^4^ both of which play a role in outflow tract development.

It is worth noting that 9 out of 11 ARIC participants who succumbed to SCD within 3 months after ECG recording were men. We cannot rule out the possibility that observed transient substrate of SCD is sex-specific. Further studies of SCD triggers in women are needed.

We observed that small SAI QRST and small SVG magnitude predicted short-term SCD, whereas large SAI QRST represented intermediate and long-term risk of SCD. Our findings are consistent with the results of the Prospective Observational Study of the ICD in Sudden Cardiac Death Prevention (PROSE-ICD), which showed that small SAI QRST was associated with increased risk of sustained ventricular tachyarrhythmias with appropriate ICD therapies during a short follow-up period^28, 29^.

### Intermediate and long-term substrates of SCD

QTc was an intermediate predictor of SCD, forecasting SCD within 6 months to 2 years. QTc reflects sympathetic activity in the ventricles of the heart^30^ which is dynamic.^31^ QTc was *not* a short-term trigger of SCD in this study. Instead, QTc characterized a short-term substrate of nonSCD, reflecting the presence of advanced structural heart disease.

All GEH measurements as well as QRS duration predicted SCD up to 15 years of follow-up. The long-term substrate of SCD was characterized by an SVG vector pointing backward (toward the LV), a wide spatial QRS-T angle, and a large SAI QRST. The peak QRS-T angle was a consistently strong long-term predictor of SCD and can be considered for life-long SCD risk prediction. Reliable long-term prediction of SCD offers an opportunity for early preventive intervention. Many GEH-associated genetic loci are implicated in cardiac development.^4^ Further studies of the underlying biology behind GEH-associated loci will help to uncover novel mechanisms of SCD and develop primary prevention strategies. A recent case-control genome-wide association study (GWAS) of sudden cardiac arrest^32^ did not identify any variants at genome-wide statistical significance. An ideal case-control study of paroxysmal life-threatening arrhythmias (e.g. SCD) would require evidence of freedom from arrhythmogenic substrate in controls, which is difficult to achieve. As both trigger and substrate are required for development of sudden cardiac arrest, a low yield from a GWAS study is to be expected. In contrast, genomic studies of electrophysiological substrates have the advantage of a more accurate measurement of phenotype and larger statistical power (as an outcome is a continuous variable), providing higher yield.

### Dynamic predictive accuracy of biomarkers within a survival framework

The dynamic nature of SCD risk is well-recognized. However, the dynamic predictive accuracy of SCD risk markers has not been previously studied. Our large prospective epidemiological study used repeated ECG measures, obtained at five follow-up visits, which ensured stable estimates of the dynamic predictive accuracy of ECG biomarkers within a survival framework. An analytical framework for the assessment of the dynamic predictive accuracy of biomarkers for censored survival data was developed fairly recently.^16^ Heagerty et al^16^ showed that a simple estimator based on the Kaplan-Meier method has serious shortcomings for characterization of accuracy for censored survival outcomes, and developed the nearest neighbor estimator as a valid ROC solution for prediction accuracy assessment which allows the censoring process to depend on the marker.

In this study, we used an analytical approach to answer an agnostic predictive accuracy question. To mimic real-life clinical scenario, we intentionally did not adjust for confounders and therefore did not comment on the independence of association of ECG biomarkers with SCD at any given time point. There were noticeable differences in the clinical characteristics of study participants who died suddenly within 3 months after ECG recording, as compared to those who experienced SCD 5 years after ECG recording. Nevertheless, observed dissimilarities in a dynamic predictive accuracy of ECG biomarkers suggested different mechanisms behind short-term SCD triggers (or transient substrates), and long-term SCD substrates. A study of SCD triggers is objectively difficult to conduct. The methodological approach of the dynamic predictive accuracy of ECG biomarkers within the survival framework can provide unique perspective on transient substrates and triggers of SCD, which prompts further investigation.

### Spatial peak vs. spatial area vectors – based GEH measurements

In vectorcardiography, there are two major approaches to define spatial vectors: either measuring spatial peak or area vectors.^33^ In our study, peak-based GEH metrics outperformed area-based GEH metrics. This finding may be at least partially explained by the fact that we used a physiologically sound definition of the heart vector origin point and time-coherent global median beat,^13^ which permitted accurate measurement of peak vectors.

### Strengths and Limitations

The strength of our study derives from the large prospective cohort design, with five longitudinal ECG recordings, long-term (median 24 years) follow-up, and a well-adjudicated SCD outcome. However, limitations of the study should be taken into account. The small number of events within 3 and 6 months after ECG recording limited statistical power of SCD trigger analyses. Replication of the SCD trigger analyses in another prospective cohort is needed. Nevertheless, this is the large prospective study of SCD triggers and substrates, suggesting differences between long-term and transient SCD-specific and non-SCD-specific substrates, reporting “expiration date” and dynamic thresholds of ECG and VCG biomarkers. In this study, correlation between GEH metrics and heart rate was weak (r values between 0.1-0.2) and we did not normalize GEH measures by heart rate. However, further studies are needed to determine whether normalization by heart rate can further improve predictive value of GEH.

## Conclusion

Dynamic predictive accuracy - an “expiration date” - of ECG and VCG biomarkers should be taken into account for development of dynamic risk scores of competing SCD risk. ECG biomarkers more accurately predicted events that occurred within the first 2 years after ECG recording, as compared to events that occurred later. Distinction between markers predicting short-term events (within 3 months) and markers predicting long-term events (15 years after ECG recording) may represent the difference between markers heralding SCD (triggers or transient substrates) versus markers identifying persistent substrate.

## Acknowledgements

The authors thank the staff and participants of the ARIC study for their important contributions. We would like to acknowledge the SCD mortality classification committee members: Nona Sotoodehnia (lead), Selcuk Adabag, Sunil Agarwal, Lin Chen, Rajat Deo, Leonard Ilkhanoff, Liviu Klein, Saman Nazarian, Ashleigh Owen, Kris Patton, and Larisa Tereshchenko.

## Funding

The ARIC study has been funded in whole or in part with Federal funds from the National Heart, Lung, and Blood Institute, National Institutes of Health, Department of Health and Human Services, under Contract nos. (HHSN268201700001I, HHSN268201700002I, HHSN268201700003I, HHSN268201700004I, HHSN268201700005I). This work was supported by HL118277 (Tereshchenko).

## Conflict of interests

None declared.

---

**Supplemental Figure 1.**
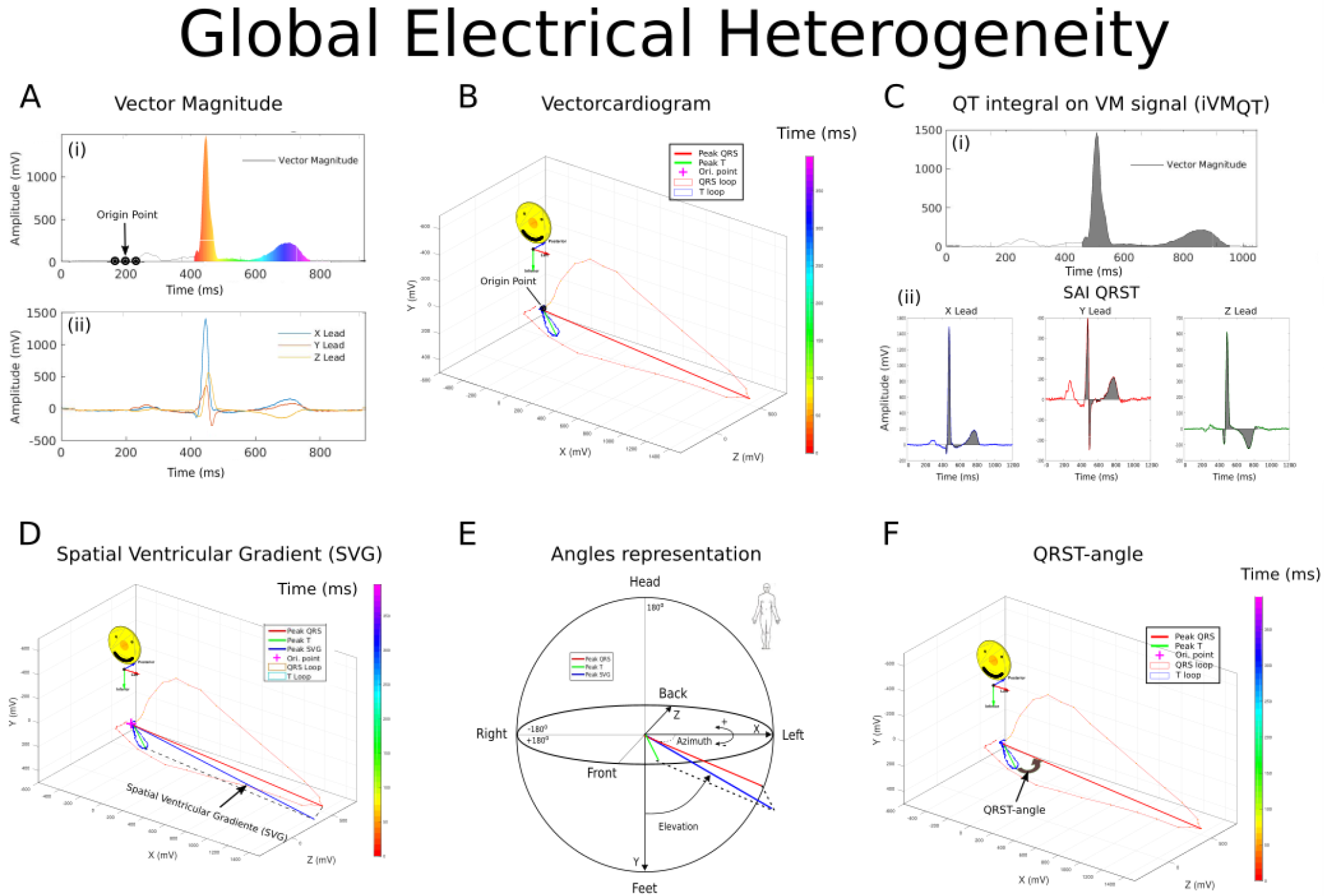
Method of measuring global electrical heterogeneity. A.(i) Vector Magnitude signal is plotted over time. Origin of the heart vector is identified as the flattest segment of the cardiac cycle, when the heart vector does not move in three-dimensional space. (ii) corresponding XYZ leads of the time-coherent median beat. B. The same Vector Magnitude is plotted in a three-dimensional space, forming vectorcardiographic loops. Origin of the heart vector is marked by a dot. C. Measurement of the scalar value of spatial ventricular gradient as (i) QT integral on Vector Magnitude signal, and (ii) Sum Absolute QRST integral (SAI QRST). D. Spatial ventricular gradient vector is a vectorial sum of spatial QRS and T vectors. Peak vectors are shown. E. Measurement of the azimuth and elevation, and magnitude of spatial ventricular gradient vector. F. Measurement of spatial peak QRS-T angle.

**Supplemental Figure 2.**
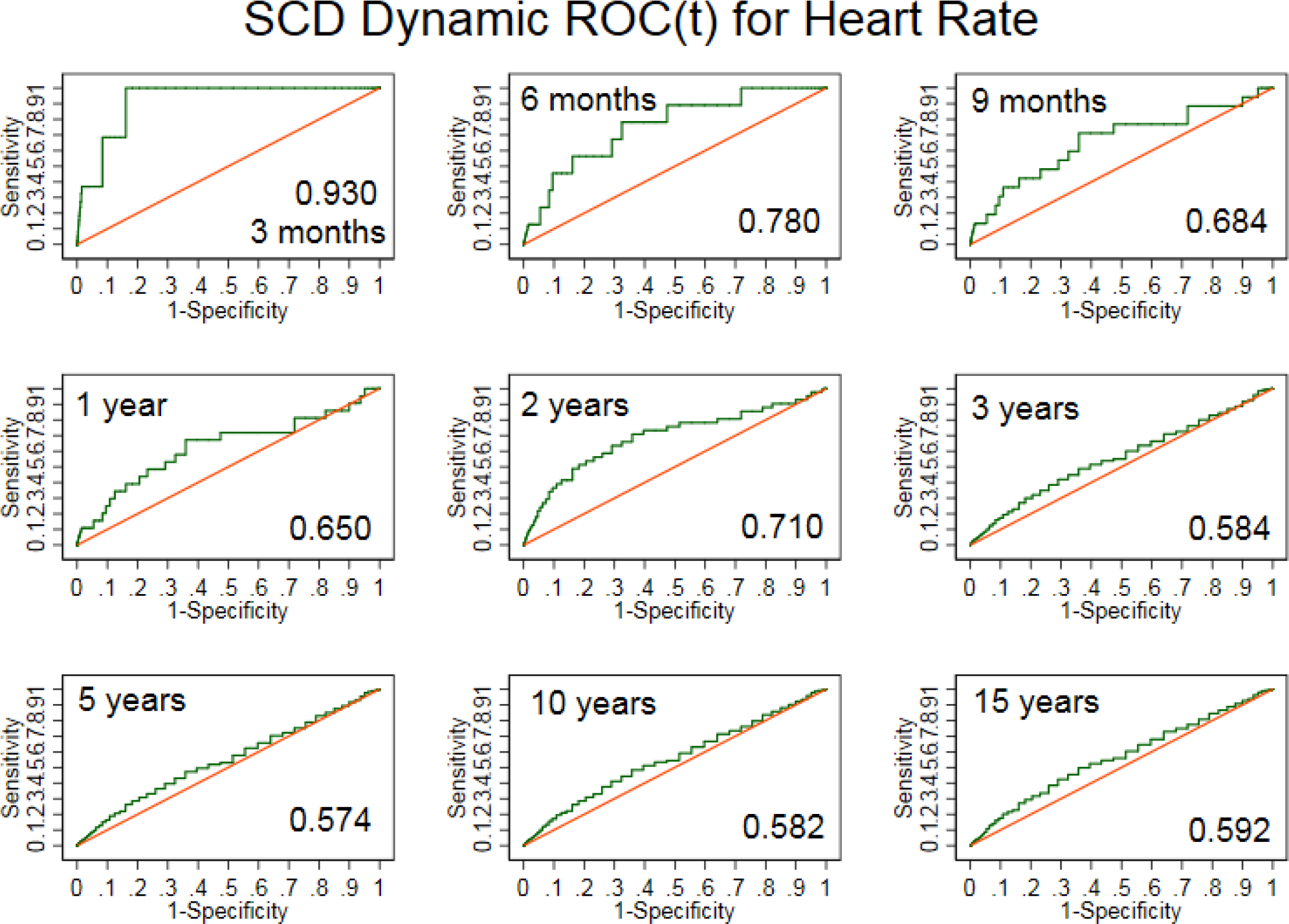
Time-dependent ROC curves for windows of sudden cardiac death prediction 3, 6, 9 months, 1, 2, 3, 5, 10, 15 years for resting heart rate.

**Supplemental Figure 3.**
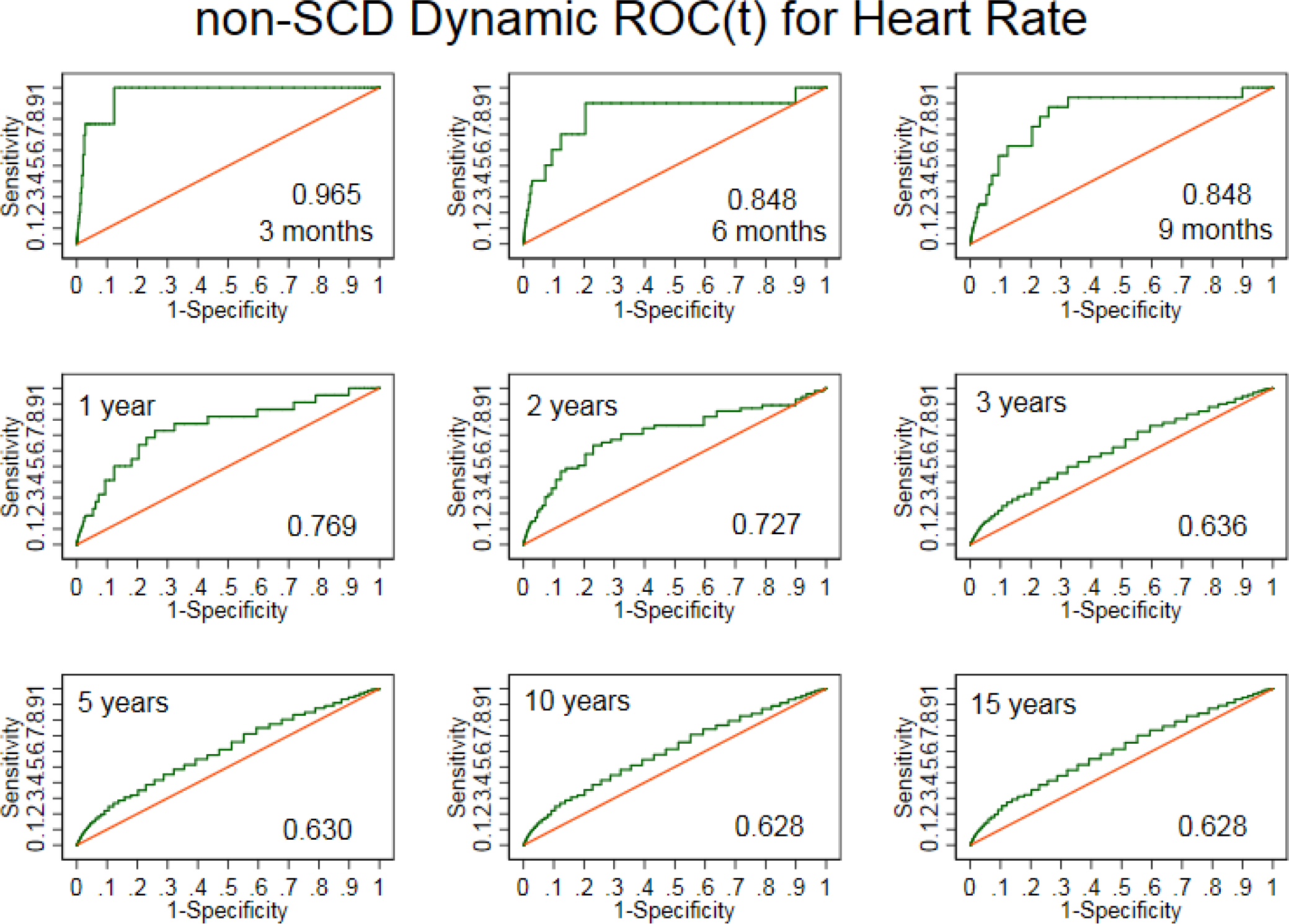
Time-dependent ROC curves for windows of non-sudden cardiac death prediction 3, 6, 9 months, 1, 2, 3, 5, 10, 15 years for resting heart rate.

**Supplemental Figure 4.**
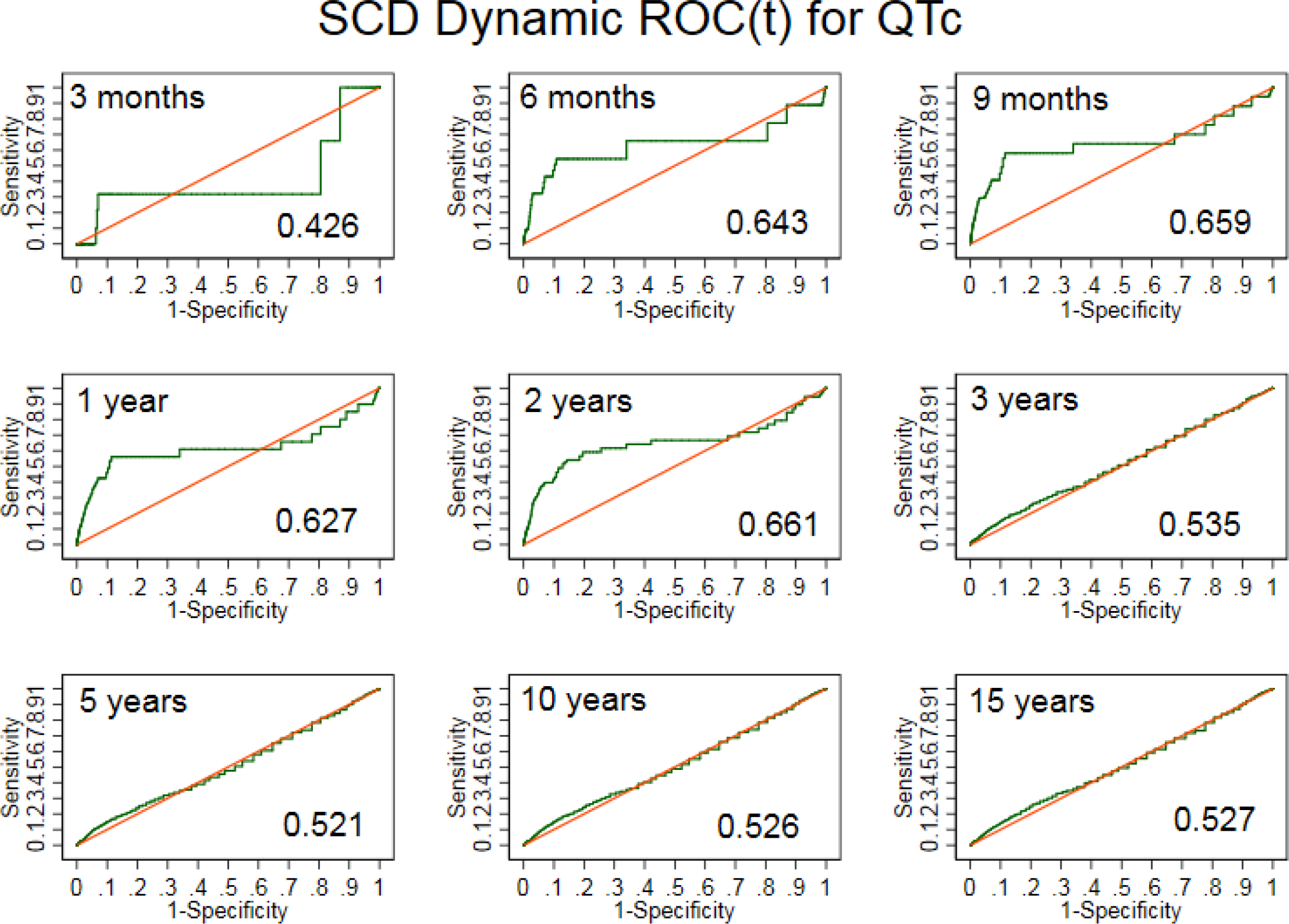
Time-dependent ROC curves for windows of sudden cardiac death prediction 3, 6, 9 months, 1, 2, 3, 5,10, 15 years for QTc interval.

**Supplemental Figure 5.**
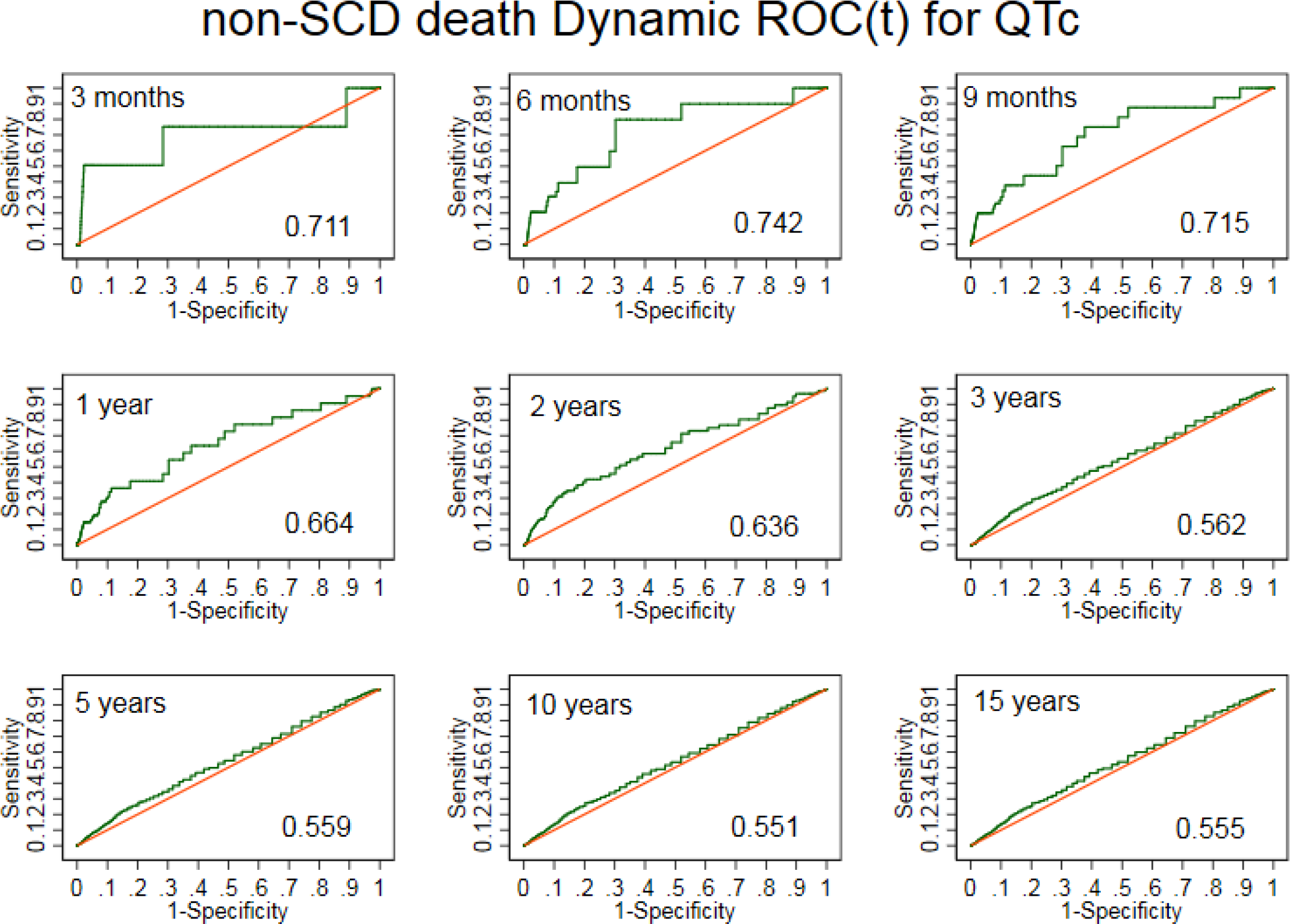
Time-dependent ROC curves for windows of non-sudden cardiac death prediction 3, 6, 9 months, 1, 2, 3, 5, 10, 15 years for QTc interval.

**Supplemental Figure 6.**
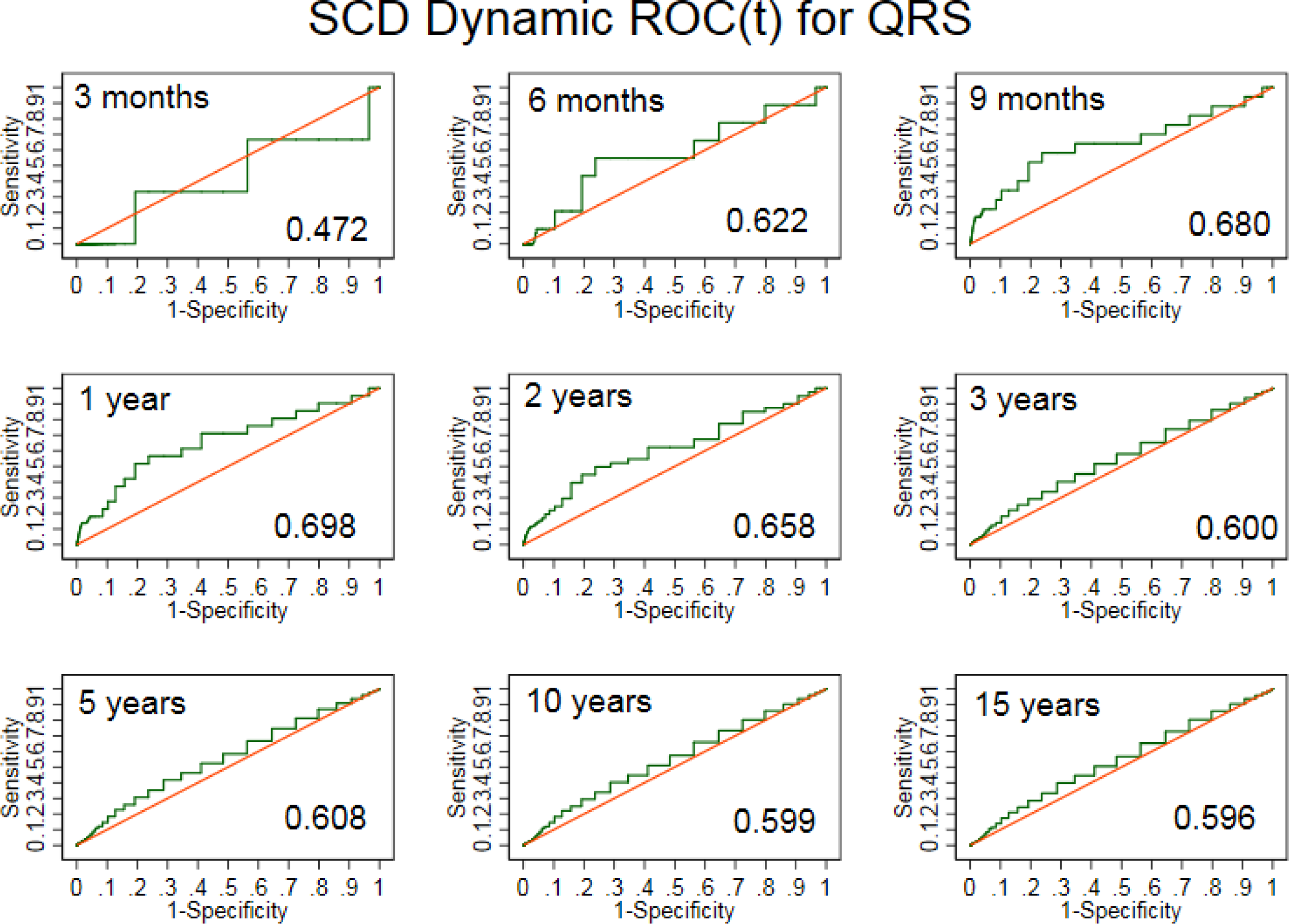
Time-dependent ROC curves for windows of sudden cardiac death prediction 3, 6, 9 months, 1, 2, 3, 5, 10, 15 years for QRS duration.

**Supplemental Figure 7.**
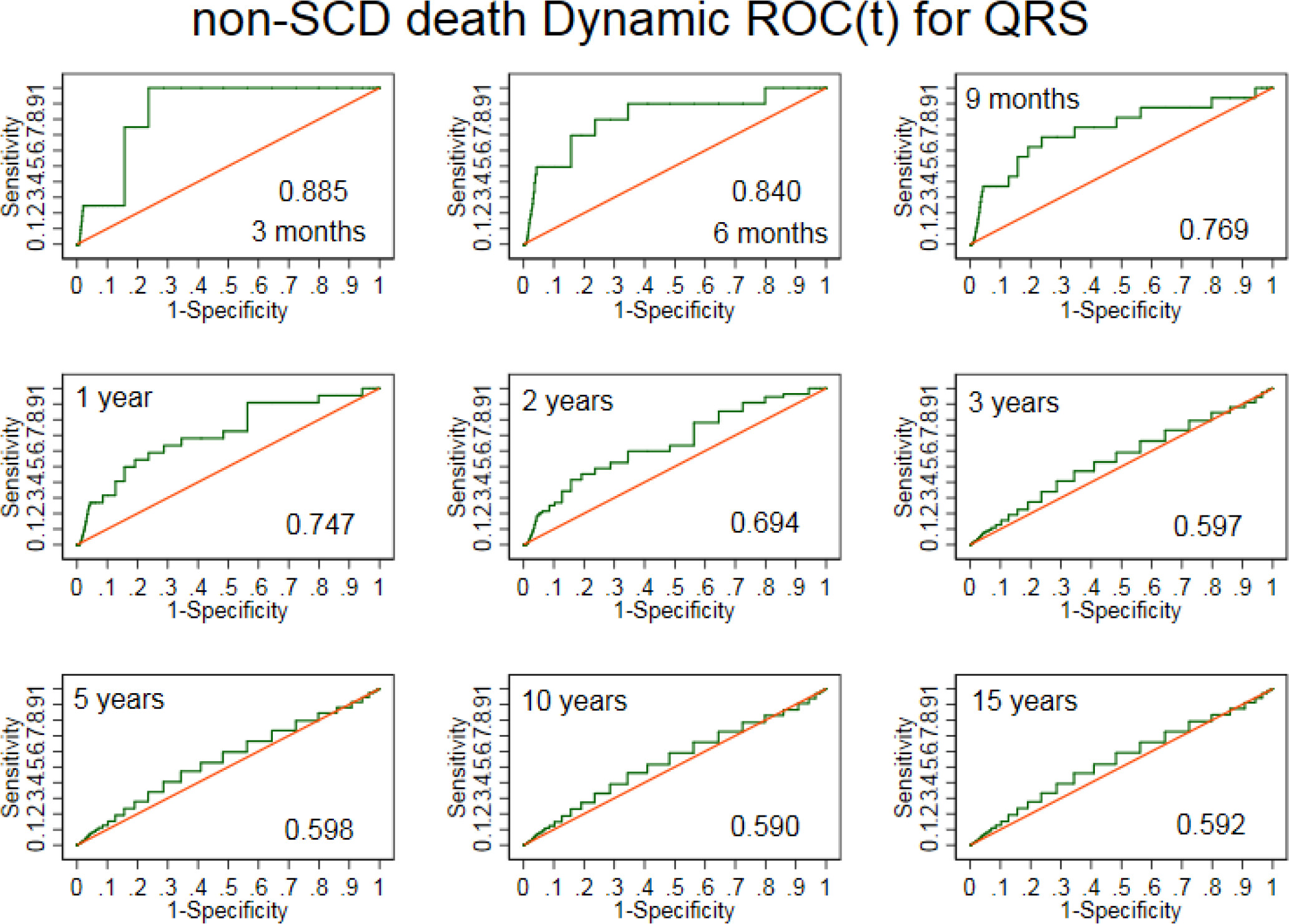
Time-dependent ROC curves for windows of non-sudden cardiac death prediction 3, 6, 9 months, 1, 2, 3, 5, 10, 15 years for QRS duration.

**Supplemental Figure 8.**
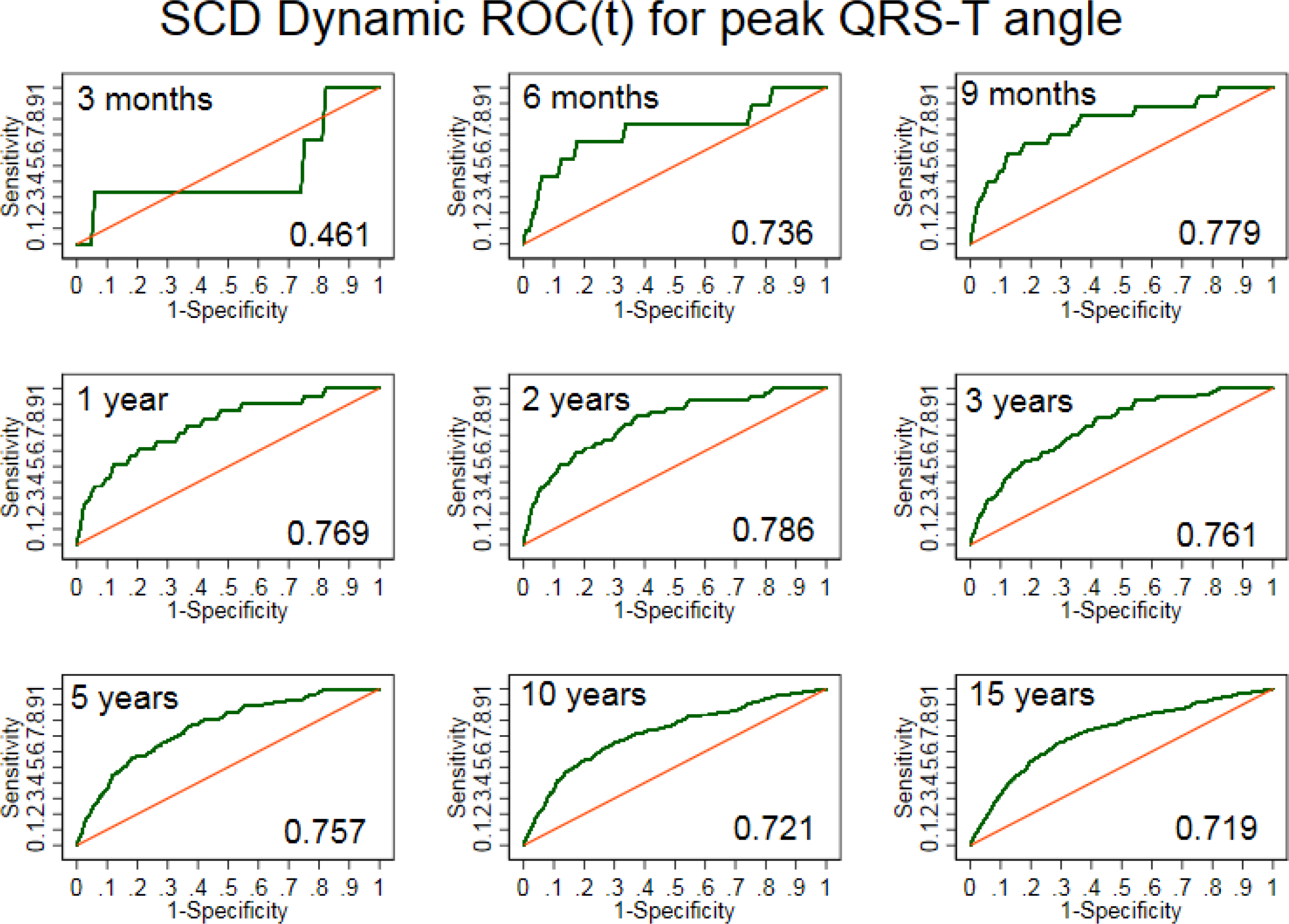
Time-dependent ROC curves for windows of sudden cardiac death prediction 3, 6, 9 months, 1, 2, 3, 5, 10, 15 years for spatial peak QRS-T angle.

**Supplemental Figure 9.**
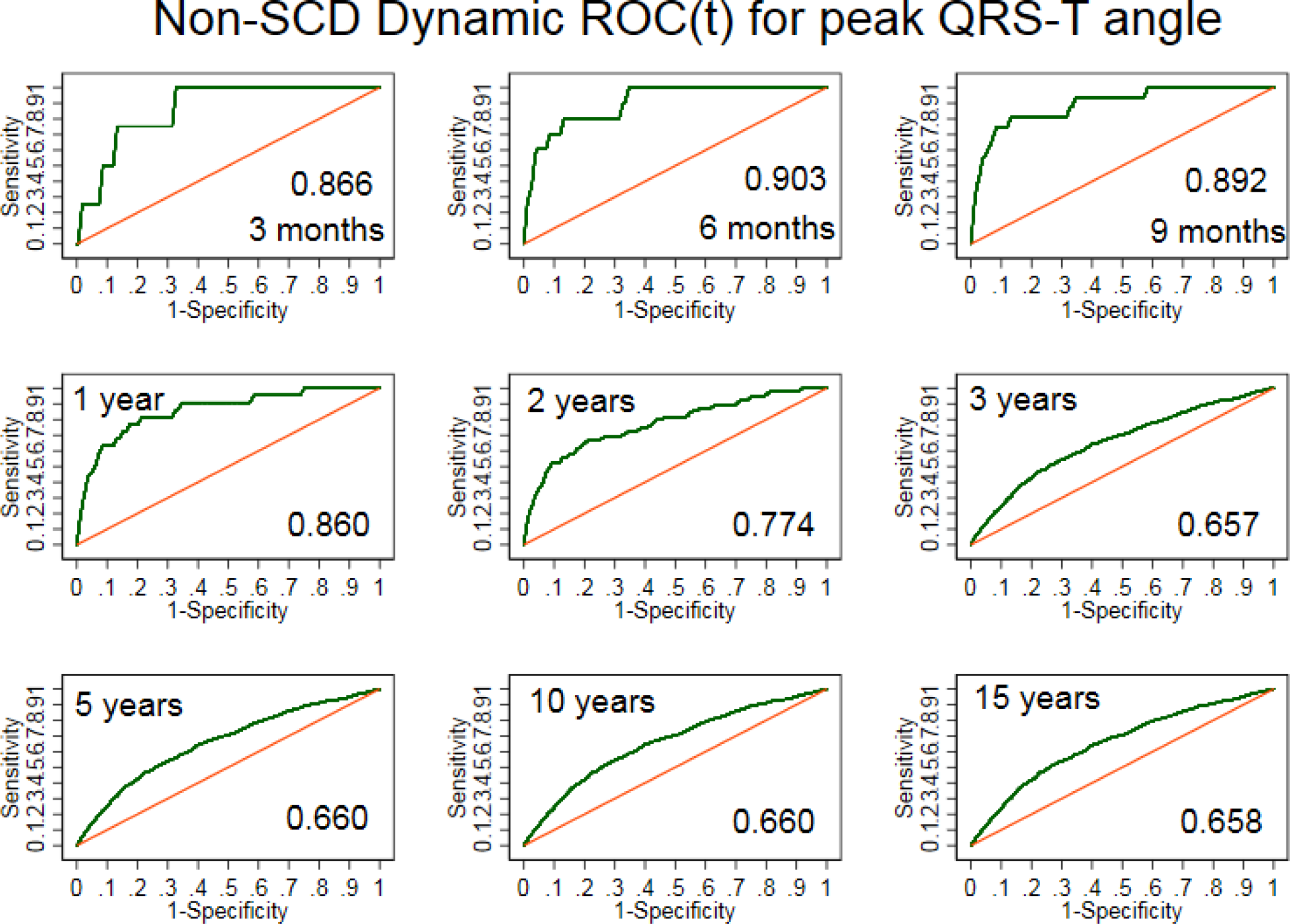
Time-dependent ROC curves for windows of non-sudden cardiac death prediction 3, 6, 9 months, 1, 2, 3, 5, 10, 15 years for spatial peak QRS-T angle.

**Supplemental Figure 10.**
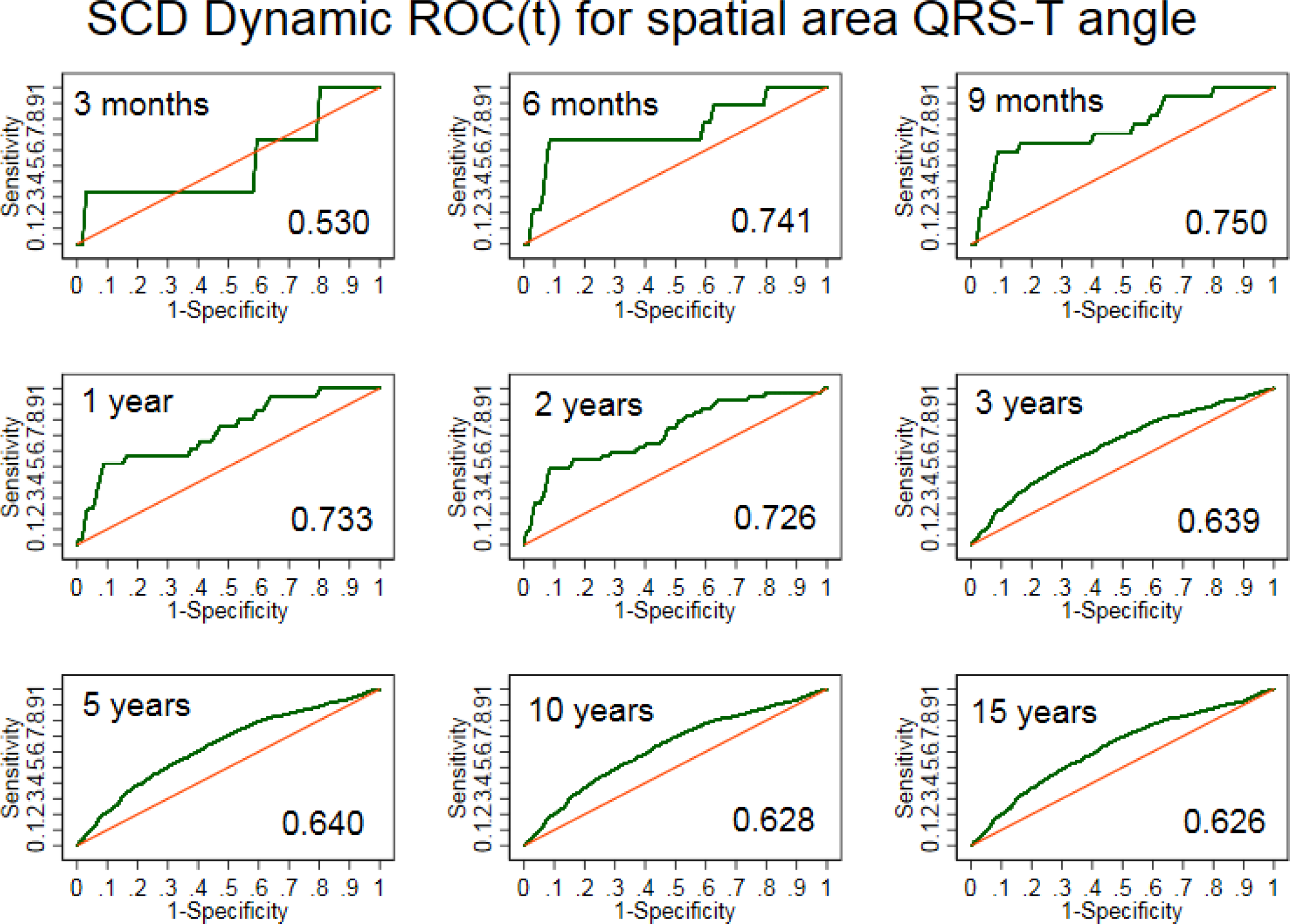
Time-dependent ROC curves for windows of sudden cardiac death prediction 3, 6, 9 months, 1, 2, 3, 5, 10, 15 years for spatial area QRS-T anglee.

**Supplemental Figure 11.**
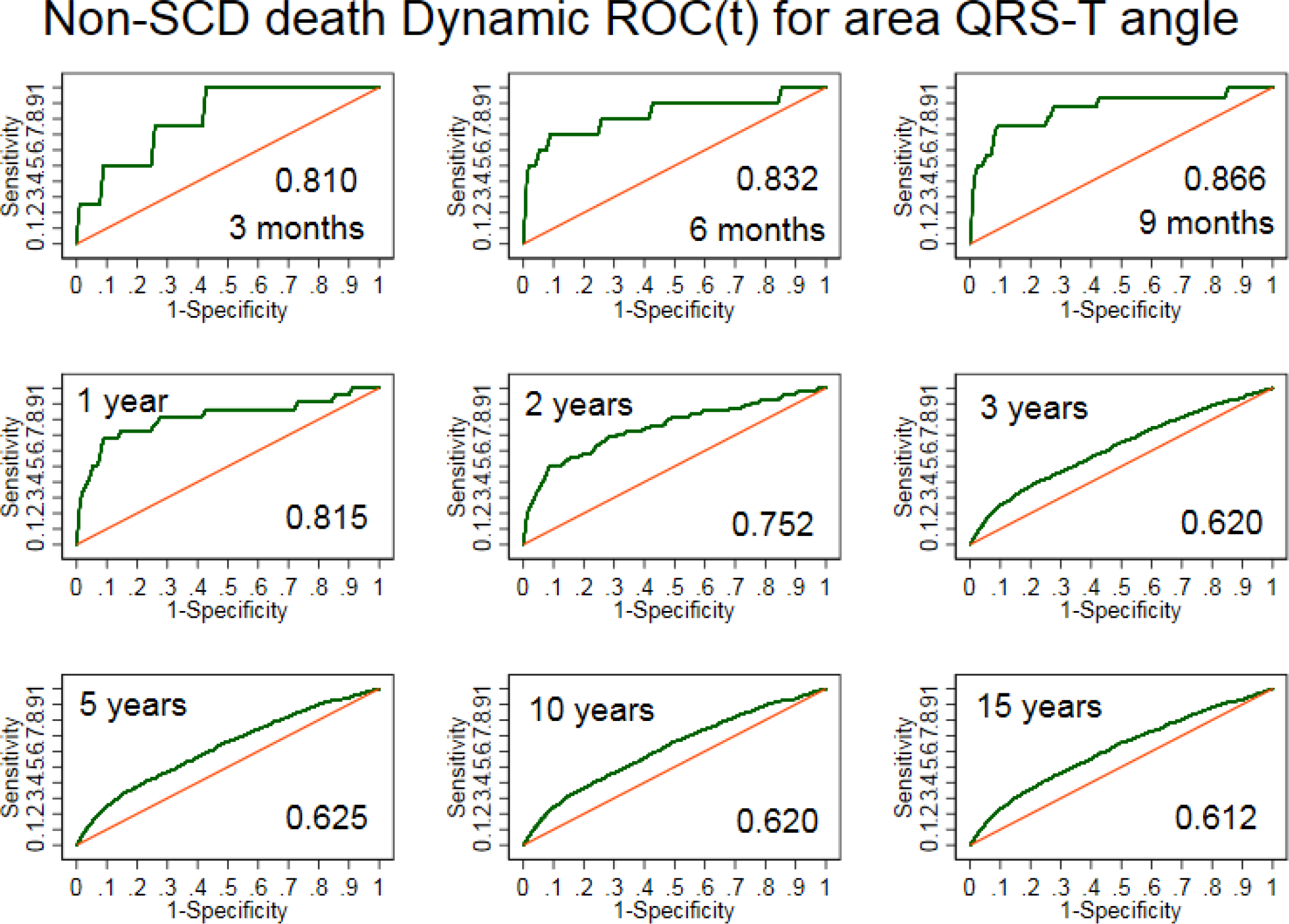
Time-dependent ROC curves for windows of sudden cardiac death prediction 3, 6, 9 months, 1, 2, 3, 5, 10, 15 years for spatial area QRS-T angle.

**Supplemental Figure 12.**
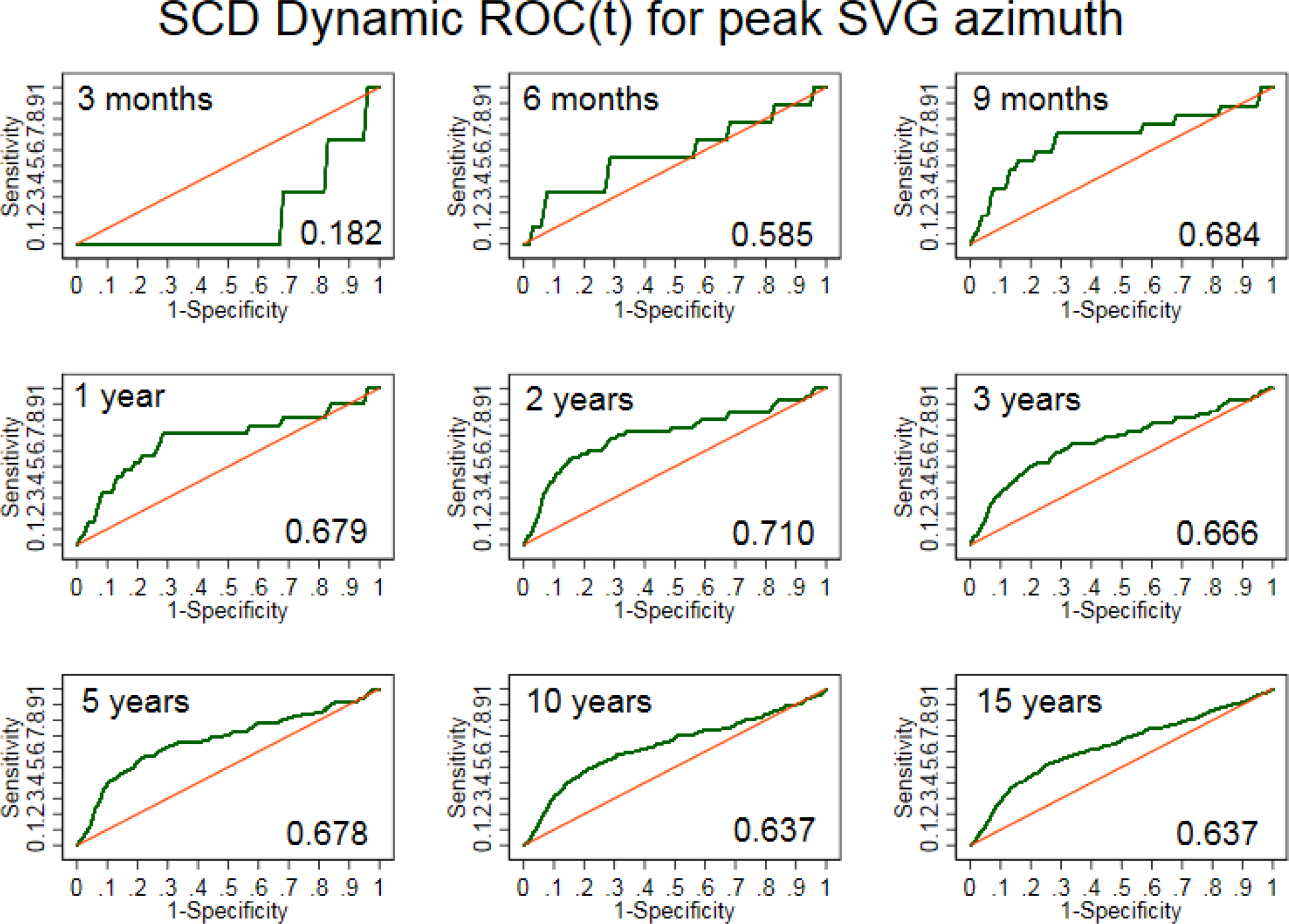
Time-dependent ROC curves for windows of sudden cardiac death prediction 3, 6, 9 months, 1, 2, 3, 5, 10, 15 years for peak spatial ventricular gradient azimuth.

**Supplemental Figure 13.**
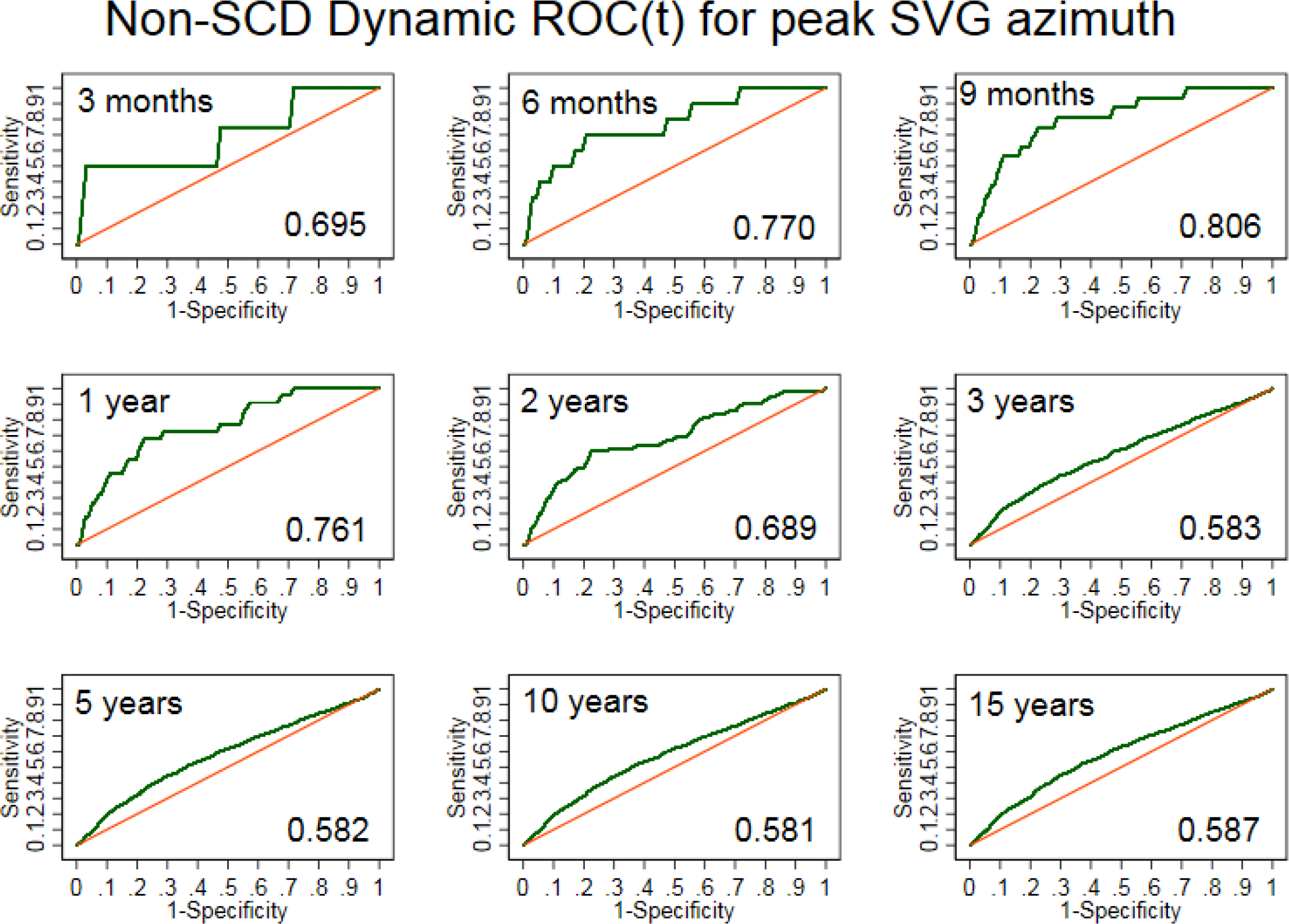
Time-dependent ROC curves for windows of non-sudden cardiac death prediction 3, 6, 9 months, 1, 2, 3, 5, 10, 15 years for peak spatial ventricular gradient azimuth.

**Supplemental Figure 14.**
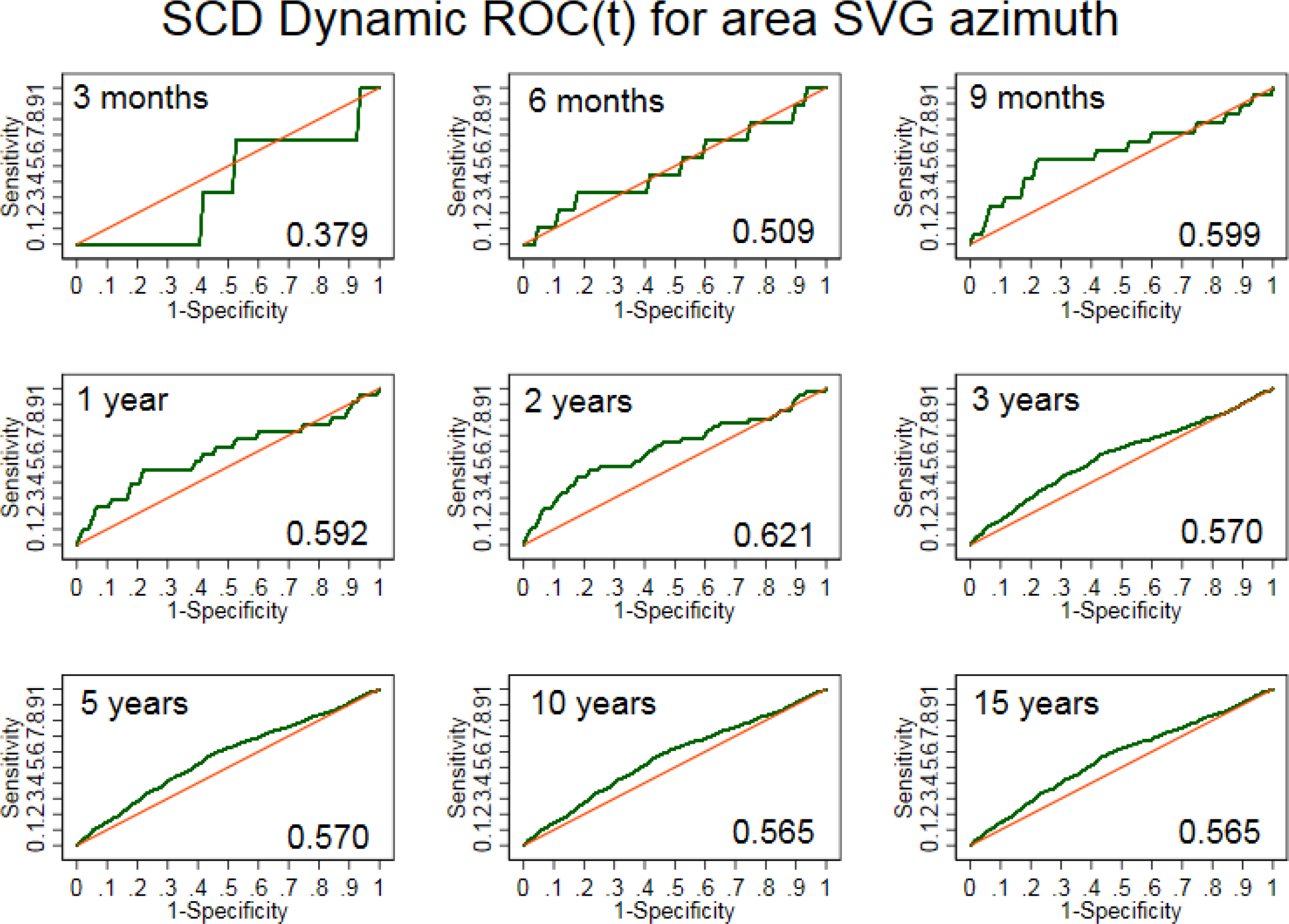
Time-dependent ROC curves for windows of sudden cardiac death prediction 3, 6, 9 months, 1, 2, 3, 5, 10, 15 years for area spatial ventricular gradient azimuth.

**Supplemental Figure 15.**
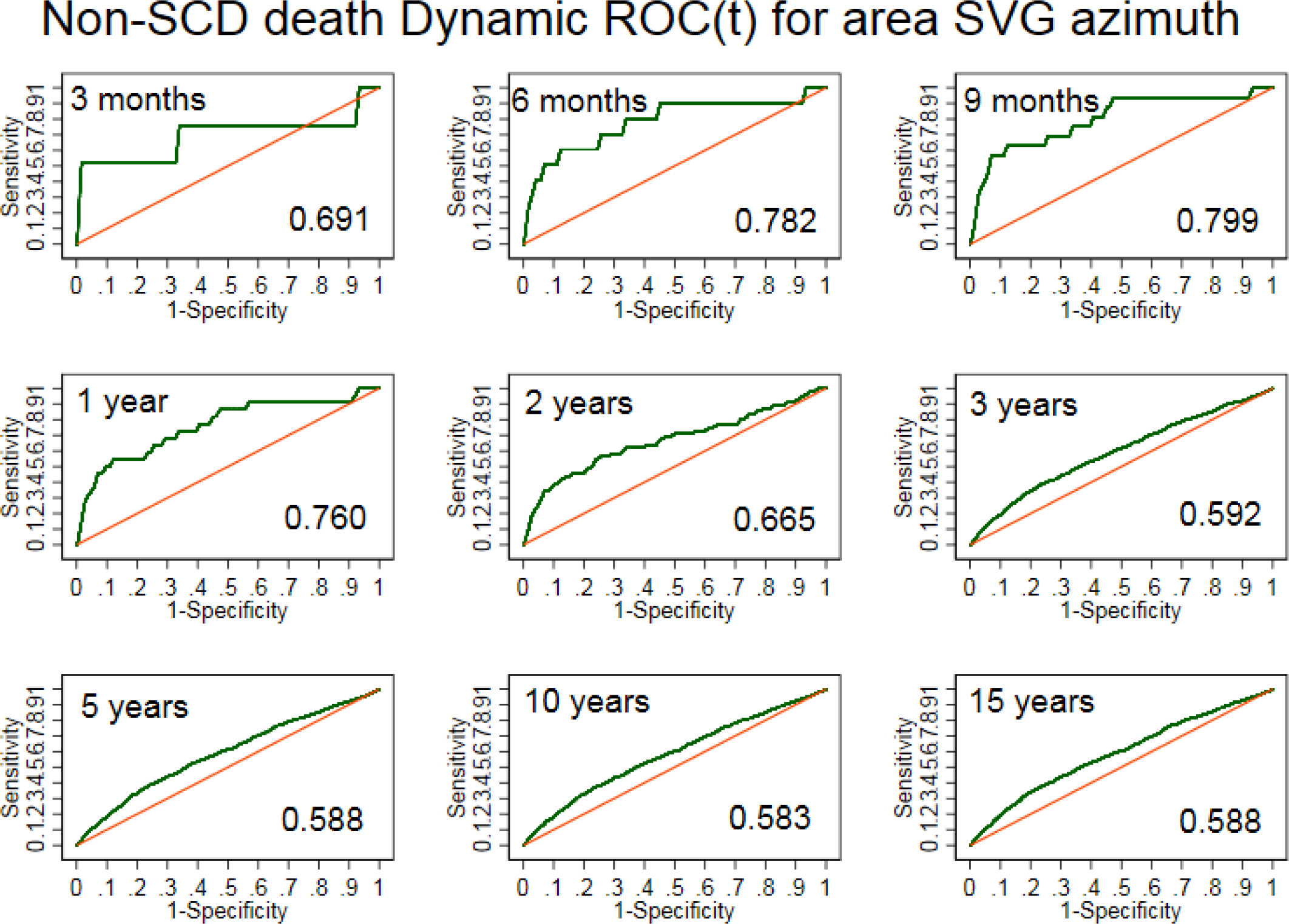
Time-dependent ROC curves for windows of non-sudden cardiac death prediction 3, 6, 9 months, 1, 2, 3, 5, 10, 15 years for area spatial ventricular gradient azimuth.

**Supplemental Figure 16.**
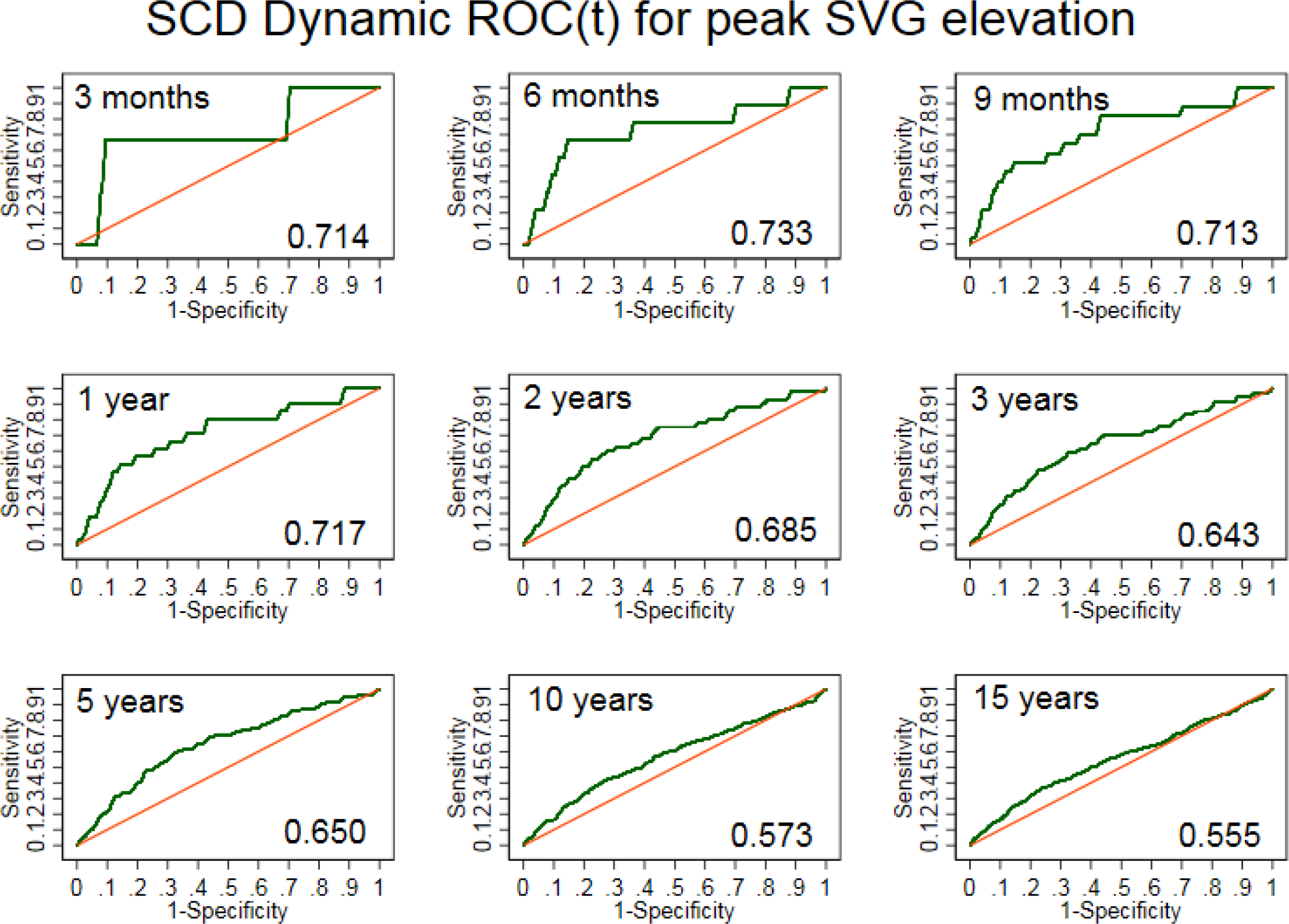
Time-dependent ROC curves for windows of sudden cardiac death prediction 3, 6, 9 months, 1, 2, 3, 5, 10, 15 years for peak spatial ventricular gradient elevation.

**Supplemental Figure 17.**
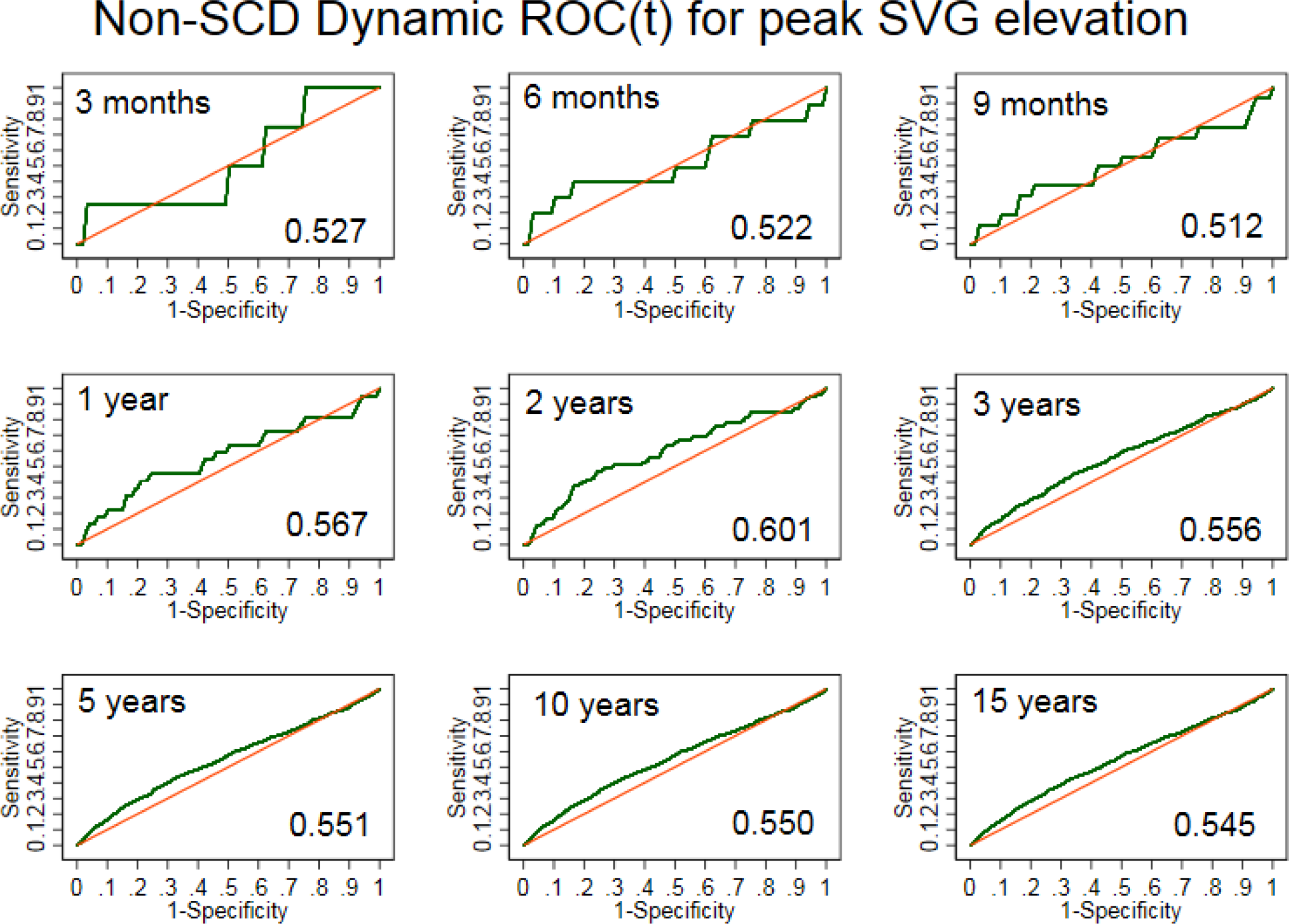
Time-dependent ROC curves for windows of non-sudden cardiac death prediction 3, 6, 9 months, 1, 2, 3, 5, 10, 15 years for peak spatial ventricular gradient elevation.

**Supplemental Figure 18.**
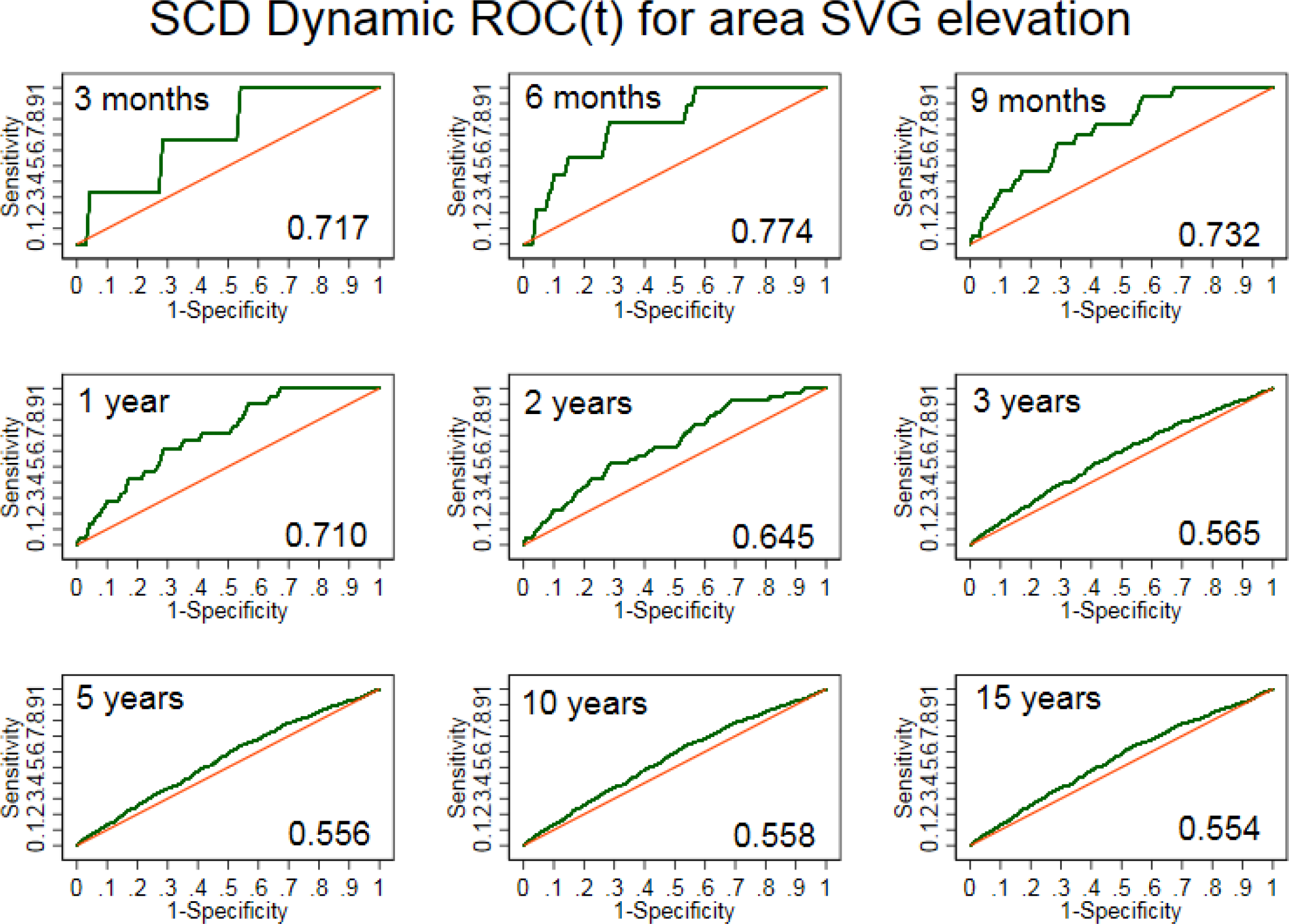
Time-dependent ROC curves for windows of sudden cardiac death prediction 3, 6, 9 months, 1, 2, 3, 5, 10, 15 years for area spatial ventricular gradient elevation.

**Supplemental Figure 19.**
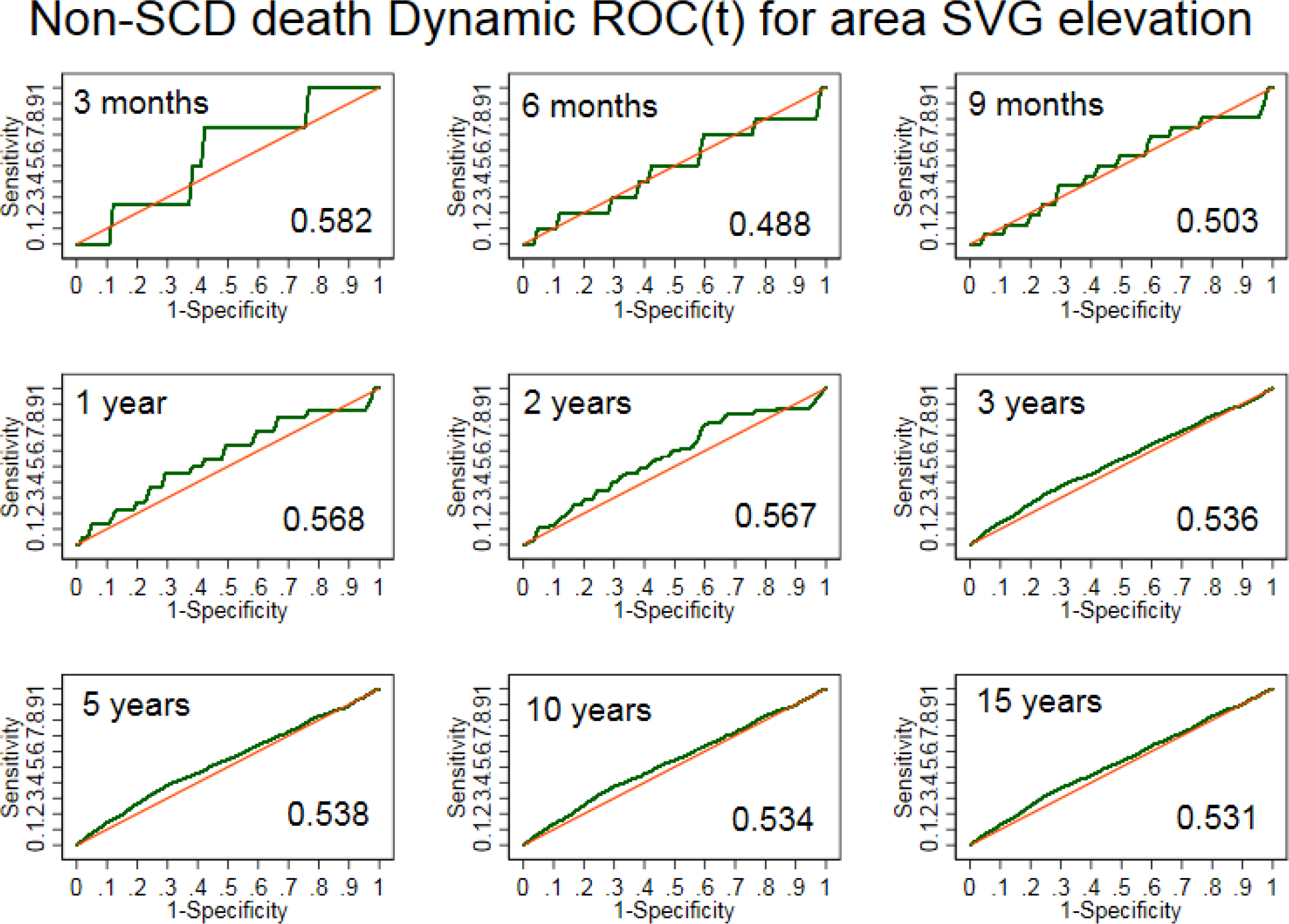
Time-dependent ROC curves for windows of non-sudden cardiac death prediction 3, 6, 9 months, 1, 2, 3, 5, 10, 15 years for area spatial ventricular gradient elevation.

**Supplemental Figure 20.**
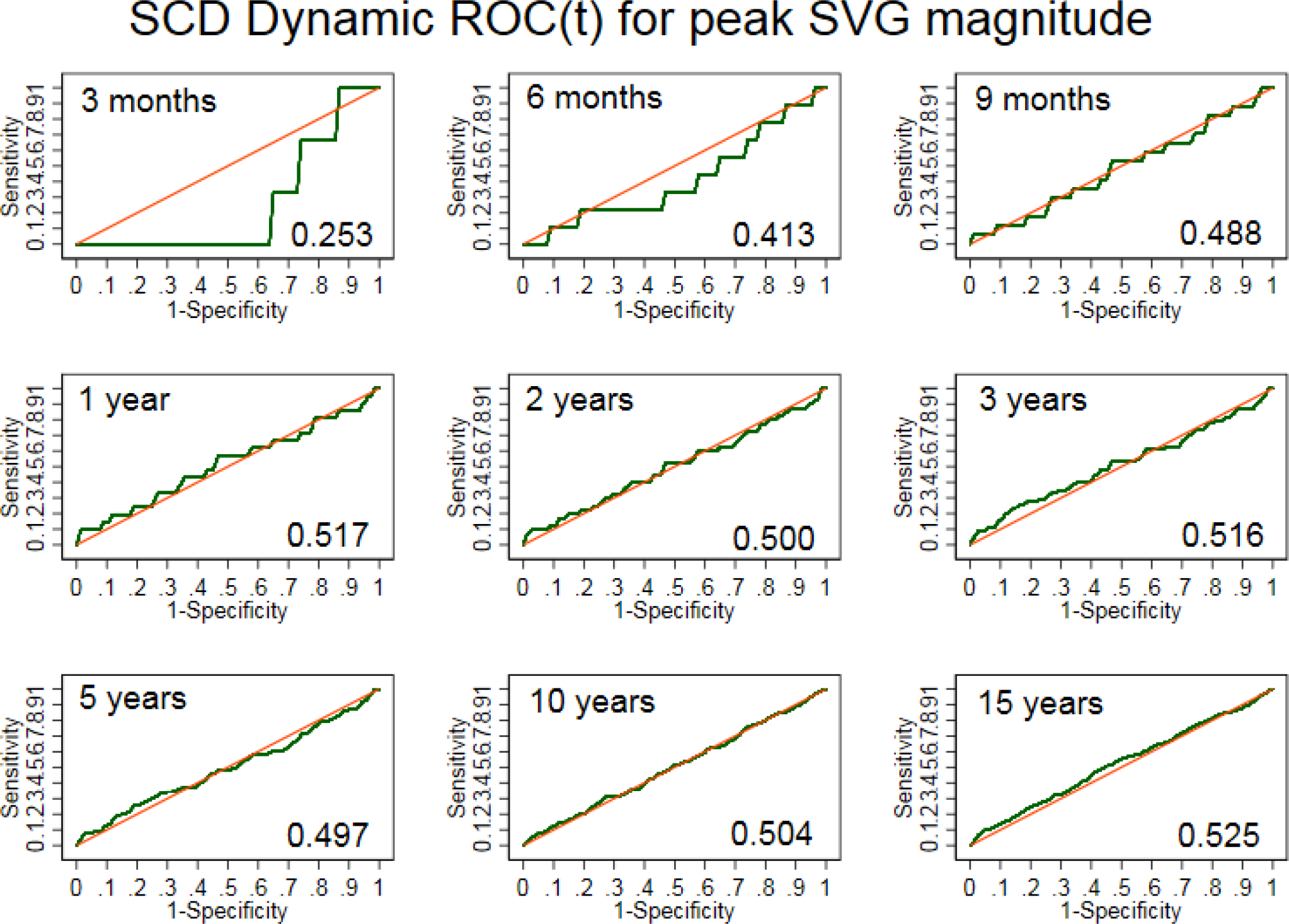
Time-dependent ROC curves for windows of sudden cardiac death prediction 3, 6, 9 months, 1, 2, 3, 5, 10, 15 years for peak spatial ventricular gradient magnitude.

**Supplemental Figure 21.**
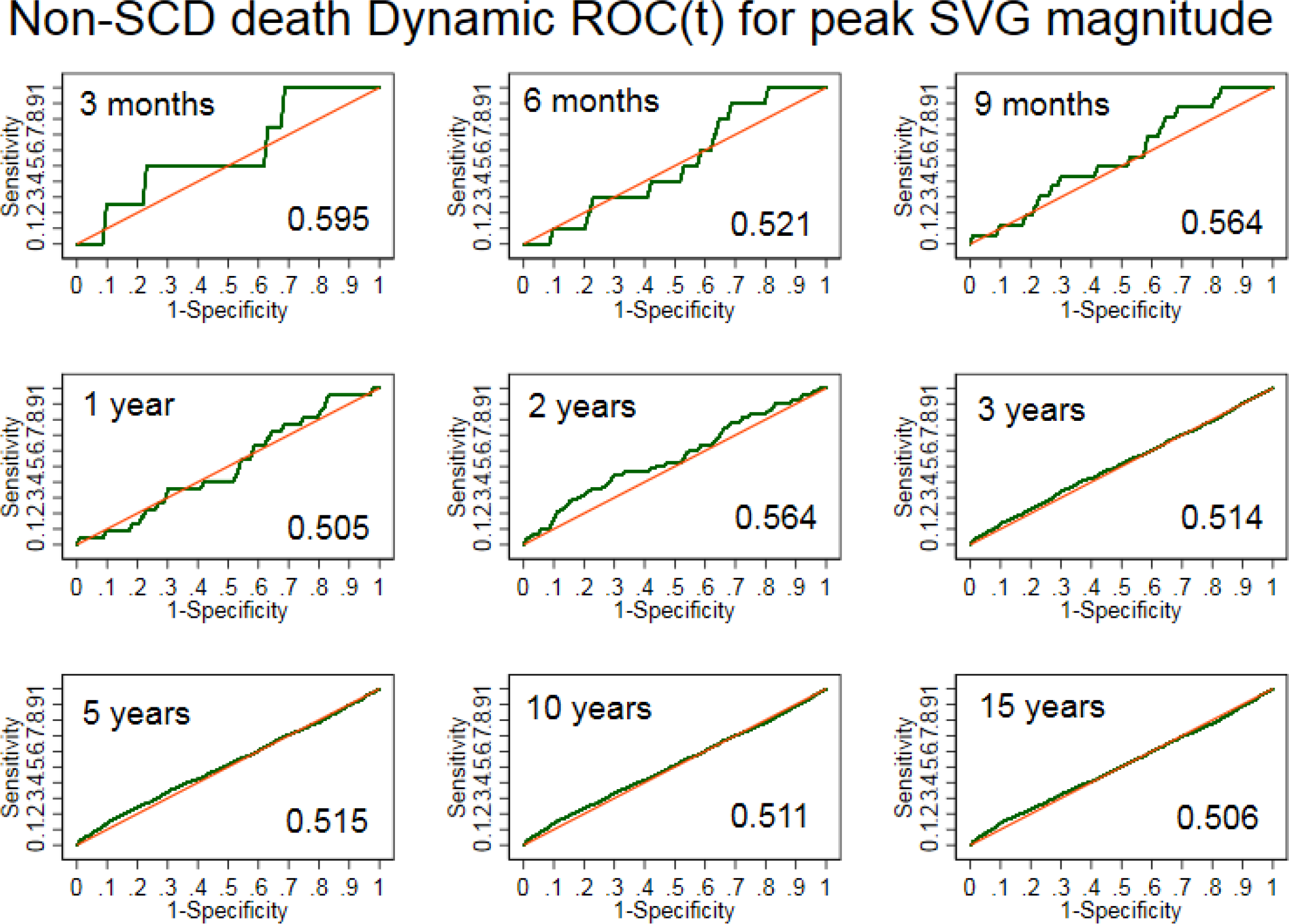
Time-dependent ROC curves for windows of non-sudden cardiac death prediction 3, 6, 9 months, 1, 2, 3, 5, 10, 15 years for peak spatial ventricular gradient magnitude.

**Supplemental Figure 22.**
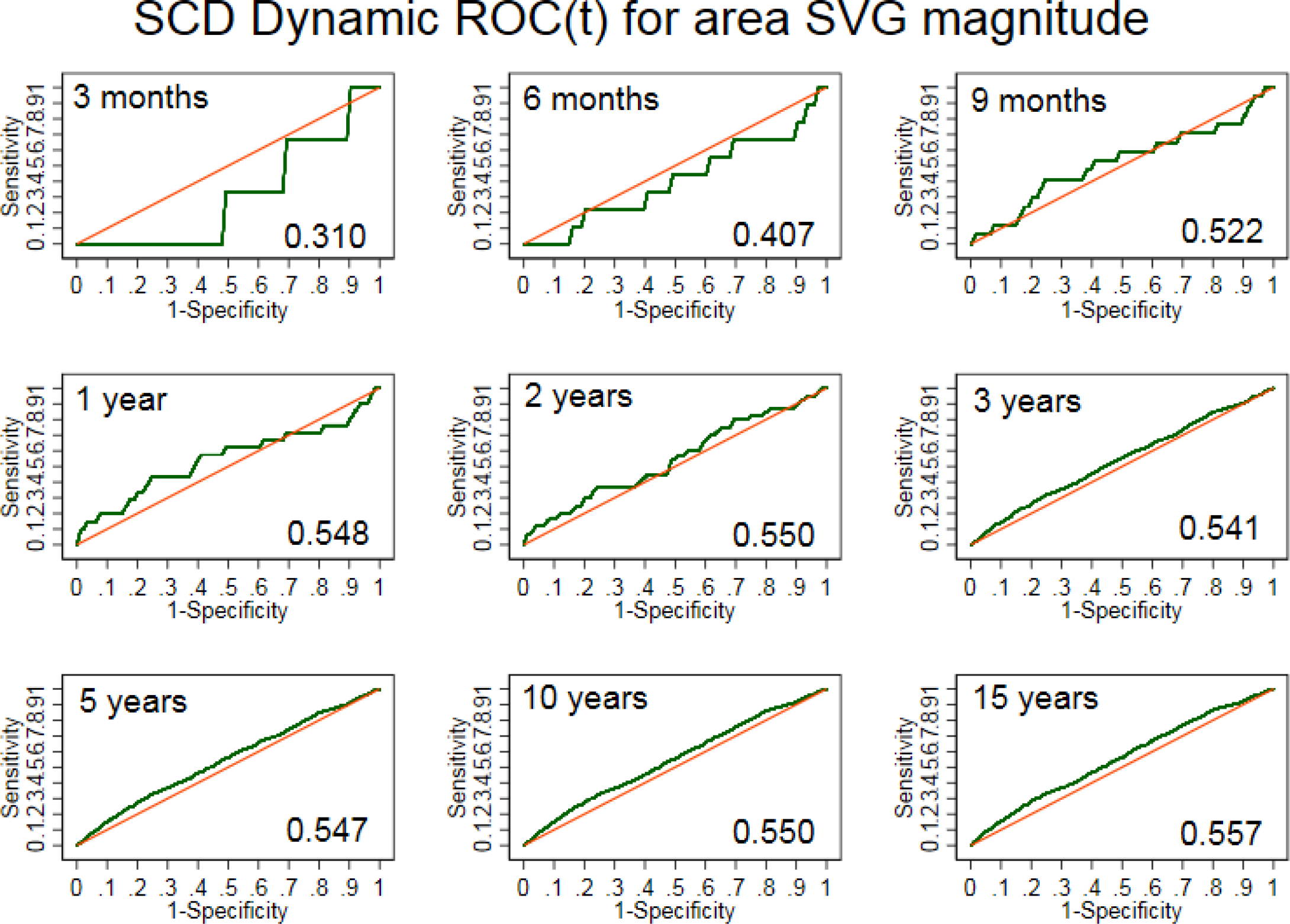
Time-dependent ROC curves for windows of sudden cardiac death prediction 3, 6, 9 months, 1, 2, 3, 5, 10, 15 years for area spatial ventricular gradient magnitude.

**Supplemental Figure 23.**
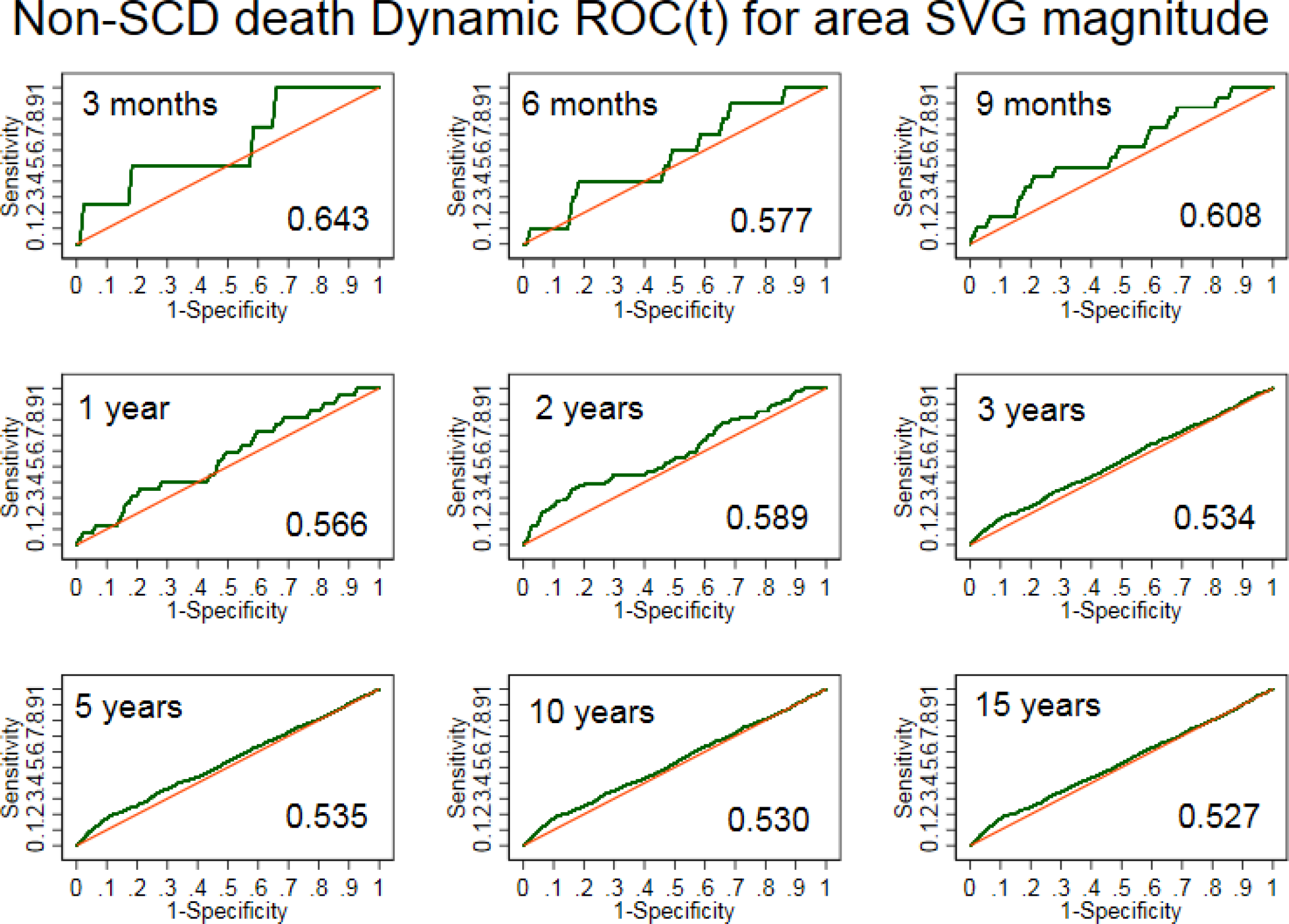
Time-dependent ROC curves for windows of non-sudden cardiac death prediction 3, 6, 9 months, 1, 2, 3, 5, 10, 15 years for area spatial ventricular gradient magnitude.

**Supplemental Figure 24.**
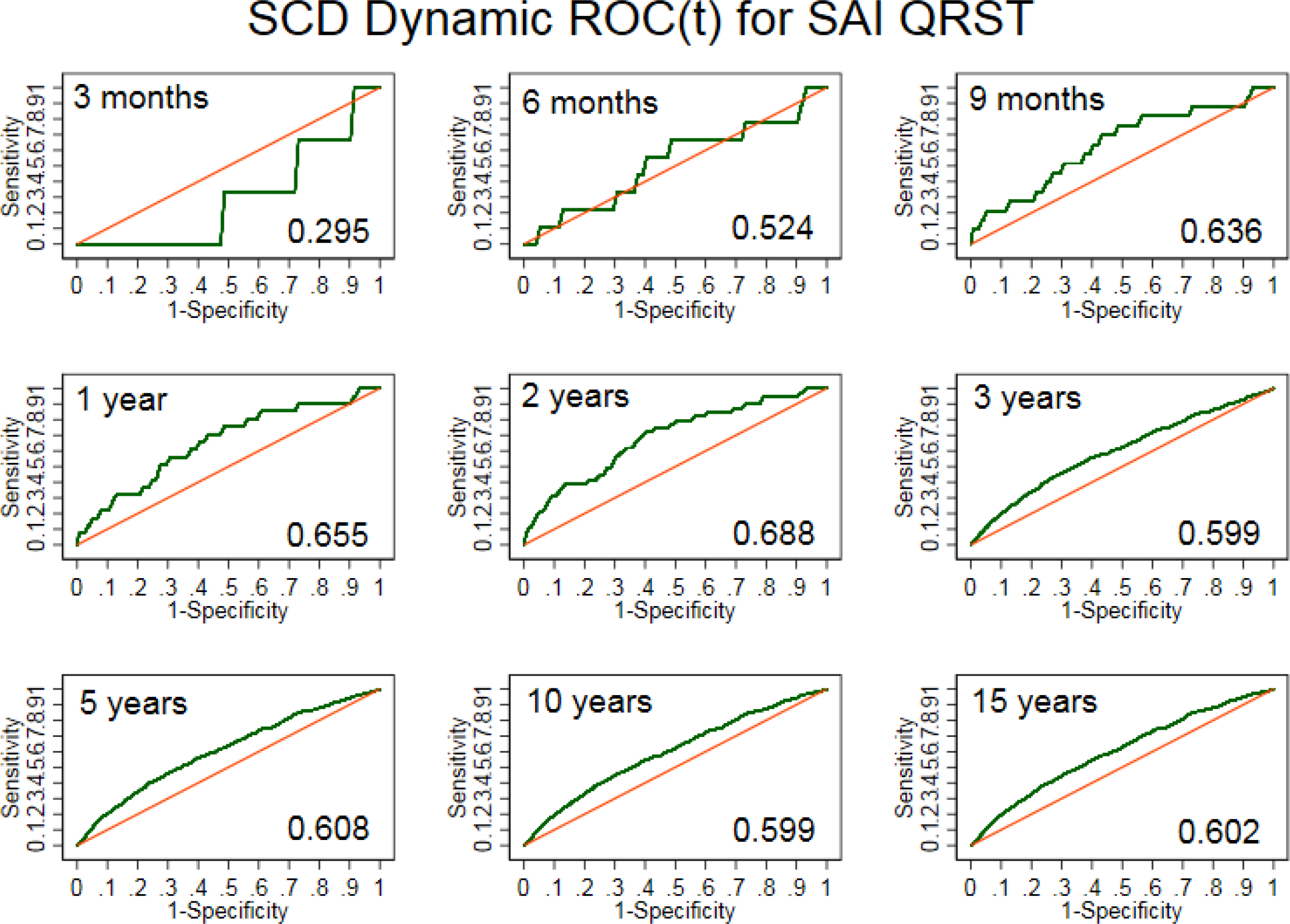
Time-dependent ROC curves for windows of sudden cardiac death prediction 3, 6, 9 months, 1, 2, 3, 5, 10, 15 years for sum absolute QRST integral (SAI QRST)

**Supplemental Figure 25.**
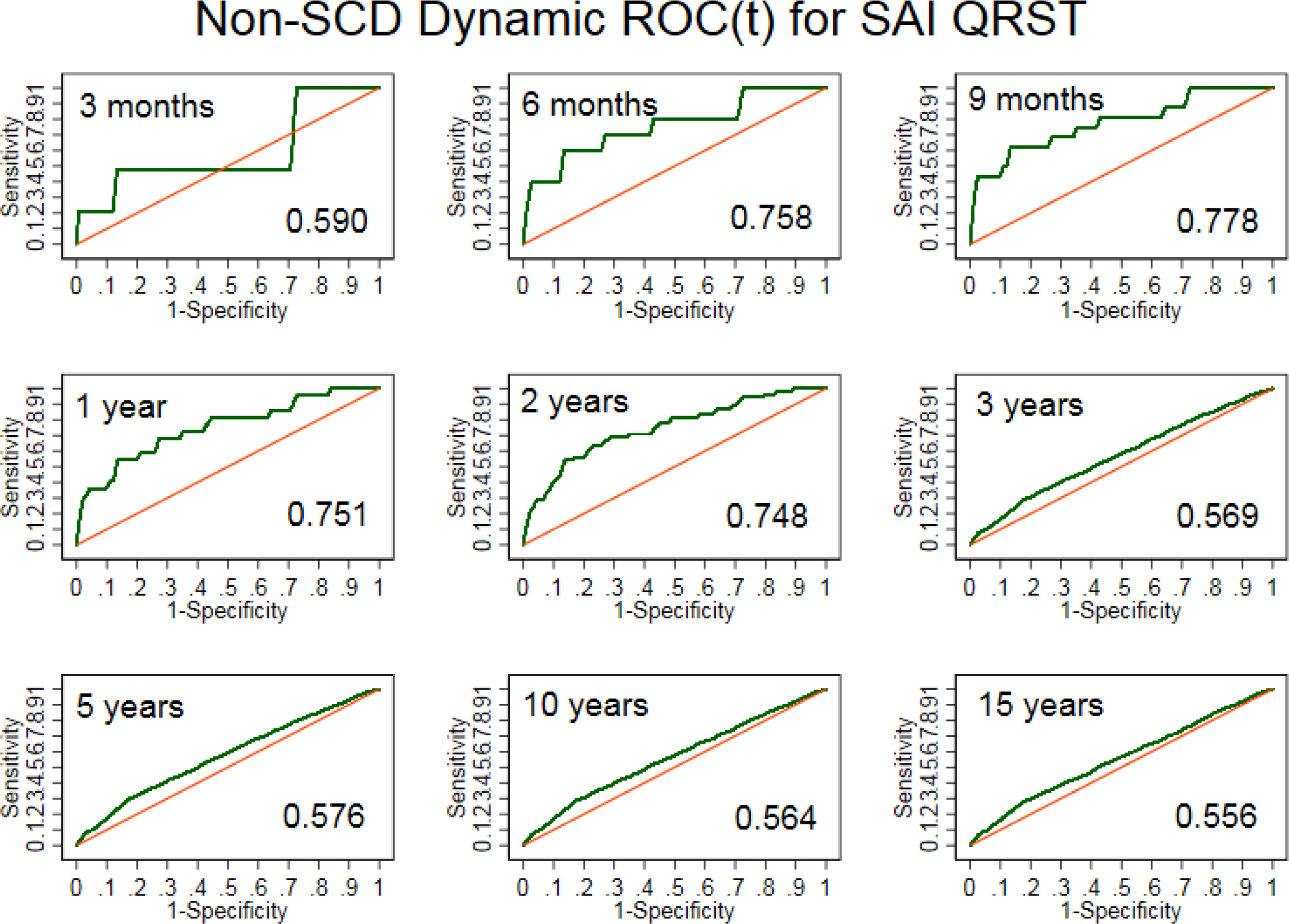
Time-dependent ROC curves for windows of non-sudden cardiac death prediction 3, 6, 9 months, 1, 2, 3, 5, 10, 15 years for sum absolute QRST integral (SAI QRST)

